# PWAS: Proteome-Wide Association Study

**DOI:** 10.1101/812289

**Authors:** Nadav Brandes, Nathan Linial, Michal Linial

## Abstract

Over the last two decades, GWAS (Genome-Wide Association Study) has become a canonical tool for exploratory genetic research, generating countless gene-phenotype associations. Despite its accomplishments, several limitations and drawbacks still hinder its success, including low statistical power and obscurity about the causality of implicated variants. We introduce PWAS (Proteome-Wide Association Study), a new method for detecting protein-coding genes associated with phenotypes through protein function alterations. PWAS aggregates the signal of all variants jointly affecting a protein-coding gene and assesses their overall impact on the protein’s function using machine-learning and probabilistic models. Subsequently, it tests whether the gene exhibits functional variability between individuals that correlates with the phenotype of interest. By collecting the genetic signal across many variants in light of their rich proteomic context, PWAS can detect subtle patterns that standard GWAS and other methods overlook. It can also capture more complex modes of heritability, including recessive inheritance. Furthermore, the discovered associations are supported by a concrete molecular model, thus reducing the gap to inferring causality. To demonstrate its applicability for a wide range of human traits, we applied PWAS on a cohort derived from the UK Biobank (~330K individuals) and evaluated it on 49 prominent phenotypes. We compared PWAS to existing methods, proving its capacity to recover causal protein-coding genes and highlighting new associations with plausible biological mechanism.

## Main

Genome-wide association studies (GWAS) seek to robustly link genetic loci with diseases and other heritable traits^1–4^. In the past decade, the method has implicated numerous variant-phenotype associations^5^ and driven important scientific discovery^6,7^. Nowadays, thanks to the rapid development of large-scale biobanks with well-genotyped and well-phenotyped cohorts, conducting case-control studies has become easier than ever. The UK Biobank (UKBB) is a flagship project of these efforts, having recruited a cohort of over 500,000 individuals, each with a full genotype and thousands of curated phenotypes (including medical history, lab tests, a variety of physical measures and comprehensive life-style questionnaires)^8,9^.

Despite the enormous impact of GWAS, inherent difficulties still limit its success^2,10^. Among the key factors are its limited statistical power, which is partly caused by the large number of tested variants across the genome. This limiting factor is especially crucial when dealing with rare variants of small effect sizes^10^. Due to Linkage Disequilibrium (LD) and population stratification, even when a genomic locus is robustly implicated with a phenotype, pinning the exact causal variant(s) is a convoluted task^6^.

Three major strategies are commonly used for prioritizing the most likely causal entities (e.g. variants or genes) affecting the phenotype. The most common strategy is fine-mapping of the raw GWAS results^11–13^. Fine-mapping of GWAS summary statistics often relies on functional annotations of the genome, under the assumption that functional entities are more likely to be causal. However, even following fine-mapping, many of the significant GWAS associations remain without any known biological mechanistic interpretation.

To arrive at more interpretable, actionable discoveries, another commonly used strategy is to prioritize genes (or other functional entities) rather than variants. There are numerous methods that aggregate GWAS summary statistics at the level of genes, often by combining them with data from expression quantitative trait locus (eQTL) studies or functional annotations of genes and pathways^14–17^.

A third strategy seeks to implicate genes directly, by carrying the association tests at the level of annotated functional elements in the first place. The most commonly used gene-level method is SKAT, which aggregates the signal across an entire genomic region, be it a gene or any other functional entity (or just a collection of SNPs)^18,19^. Another approach, recently explored by methods such as PrediXcan^20^ and TWAS^21^, tests whether the studied phenotypes correlate with gene expression levels predicted from genetic variants. Under this paradigm, the association test is comprised of two stages. First, an independent reference panel is used to train a prediction model of gene expression (in a particular tissue) as a function of the genetic makeup of a sample. The learned model is then applied on the phenotyped dataset, and the predicted gene expression levels are tested against phenotypes of interest. The advantages of this approach include a reduced burden of multiple testing, as well as more concrete and interpretable discoveries.

A natural enhancement to these approaches would be a protein-centric method that considers the effects of genetic variants on the functionality of genes, rather than affecting their abundance (be it at the transcript or protein level).

We present PWAS: Proteome Wide Association Study (Fig. 1). PWAS is based on the premise that causal variants in coding regions affect phenotypes by altering the biochemical functions of the genes’ protein products (Fig. 1a). Such functional alterations could be, for example, changes to a protein’s enzymatic activity or binding capacity (e.g. of a ligand, DNA/RNA molecule, or another protein). To capture these effects, PWAS quantifies the extent to which proteins are damaged given an individual’s genotype. Specifically, PWAS considers any variant that affects the coding-regions of genes (e.g. missense, nonsense, frameshift). It quantifies the impact of these variants on the function of the affected proteins using FIRM, a machine-learning model that considers the rich proteomic context each affecting variant^22^. These predicted effects are then combined with the genotyping data of the cohort and aggregated into per-gene functional predictions, where each protein-coding gene is assigned functional effect scores (Fig. 1b). For each gene (in the context of a specific individual) PWAS assigns two scores, to cover the two elementary modes of heritability: dominant and recessive inheritance (other modes of heritability can also be represented as a composition of the two). Intuitively, the dominant effect score is intended to express the probability of at least one hit damaging the protein function, while the recessive score attempts to express the probability of at least two damaging hits. PWAS then tests, using routine statistical analysis, if a gene’s effect scores are associated with the phenotype. In the case of a binary phenotype, a significant correlation would mean that the effect scores of cases are different than those of controls, namely that the protein is more (or less) damaged in affected individuals.

**Fig. 1:**
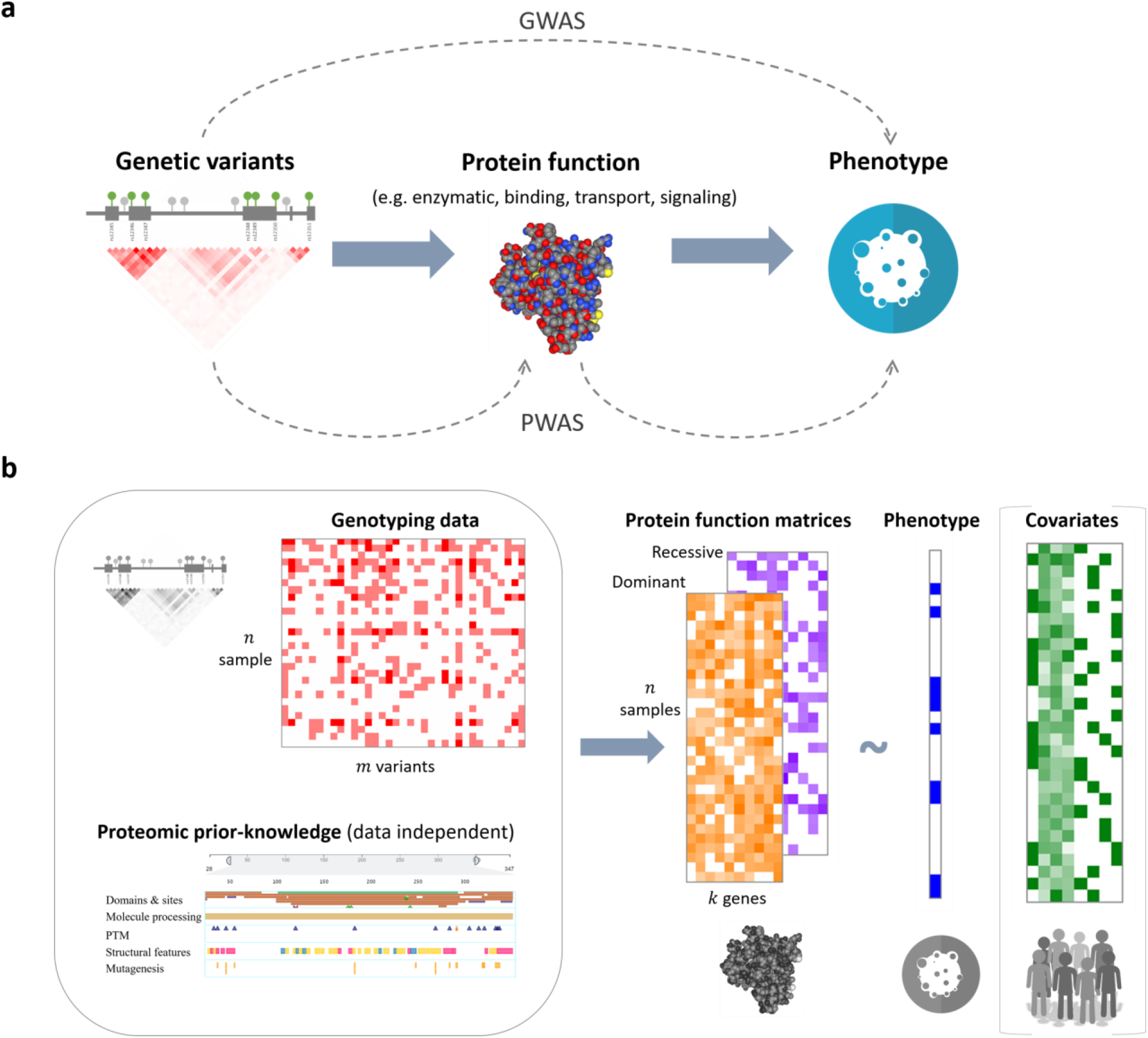
The PWAS framework. (**a**) The causal model that PWAS attempts to capture: genetic variants (within a coding region) affect the function of a protein, whose altered function influences a phenotype. PWAS identifies protein-coding genes whose overall genetic functional alterations are associated with the studied phenotype by explicitly modeling and quantifying those functional alterations. In contrast, GWAS seeks direct associations between individual variants and the phenotype. (**b**) Overview of the PWAS framework. PWAS takes the same inputs as GWAS: i) called genotypes of *m* variants across *n* individuals, ii) a vector of *n* phenotype values (could be either binary or continuous), and iii) a covariate matrix for the *n* individuals (e.g. sex, age, principal components, batch). By exploiting a rich proteomic knowledgebase, a pre-trained machine-learning model estimates the extent of damage caused to each of the *k* proteins in the human proteome, as a result of the *m* observed variants, for each of the *n* individuals (typically *k* << *m*). These estimations are stored as protein function effect score matrices. PWAS generates two such matrices, reflecting either a dominant or a recessive effect on phenotypes. PWAS identifies significant associations between the phenotype values to the effect score values in the columns of the matrices (where each column represents a distinct protein-coding gene), while taking into account the provided covariates. Each gene can be tested by the dominant model, the recessive model, or a generalized model that uses both the dominant and recessive values.

Like other gene-based approaches, PWAS enjoys a reduced burden for multiple-testing correction. In addition, it provides concrete functional interpretations for the protein-coding genes it discovers (Fig. 1a). By aggregating the signal spread across all the variants affecting the same gene, it can uncover associations that would remain undetectable at per-variant resolution, especially when rare variants are involved.

To examine the properties of PWAS, we first test it on simulated data, analyzing its statistical power across different settings. We then test it on real data derived from the UKBB, to demonstrate its wide applicability across a diverse set of phenotypes. We further compare the results of PWAS to established methods, specifically to standard GWAS and SKAT. Finally, we highlight associations uniquely discovered by PWAS.

## Results

### Functional effect scores

We analyzed a cohort derived from the UKBB. Of ~18K analyzed protein-coding genes, 17,843 were affected by at least one non-synonymous variant reported in the UKBB. On average, each of these genes was affected by 35.9 such variants (Fig. 2a).

**Fig. 2:**
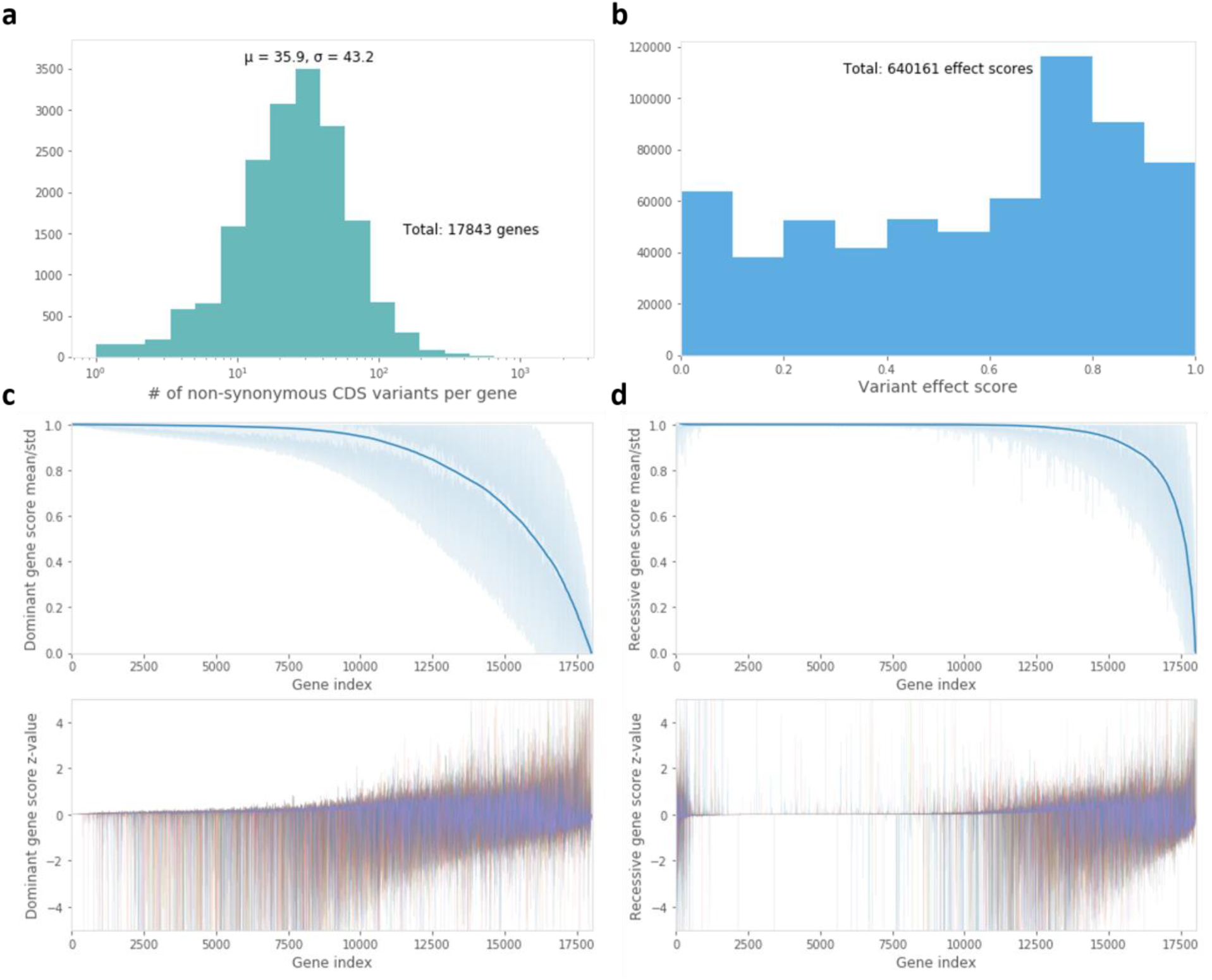
Predicted genetic functional effect scores in the UKBB cohort. (**a**) The distribution of the number of non-synonymous variants per gene that affect its coding region (in log scale), according to the (imputed) genetic data of the UKBB. (**b**) The distribution of the ~640K variant effect scores. Each score is a number between 0 (complete loss of function) to 1 (no damage to the protein product). (**c-d**) Aggregated gene scores according to the dominant (c) and recessive (d) inheritance models. Top panels: the mean (solid line) and standard deviation (shaded area) of the effect scores of the 18,053 analyzed protein-coding genes across the entire UKBB cohort (sorted by the mean score). Bottom panel: z-values of the gene effect scores across 10 randomly selected samples (of the entire ~500K samples in the UKBB). Each of the 10 samples is shown in a distinct color.

The derivation of the gene effect score matrices is comprised of two steps. First, FIRM is used to predict an effect score for each protein-affecting variant (Fig. 2b). Intuitively, these predicted effect scores can be interpreted as the probability of the variant-affected protein to retain its function. The variant scores are then integrated with the cohort genotypes and aggregated together to derive per-sample dominant and recessive effect scores at the gene level (Fig. 2c-d). As expected, dominant genetic effects (capturing single hits) are more prevalent than recessive effects (of double hits). The derived gene scores capture genetic variability in the UKBB population observed even within a small number of samples. The objective of PWAS is to test whether this functional genetic variability correlates with phenotypes.

### Simulation analysis

To examine the discovery potential of PWAS compared to GWAS and SKAT, we conducted a simulation analysis (Fig. 3). The simulation was carried on real genetic data (from the UKBB cohort), with phenotypes simulated by mixing genetic signal and noise. To test the sensitivity of PWAS to the inevitable inaccuracies of FIRM, we examined the effect of a noise parameter (*ϵ*) influencing its predictions. It appears that under the modeling assumptions of the simulation, PWAS is not very sensitive to limited inaccuracies of the underlying machine-learning predictor.

**Fig. 3:**
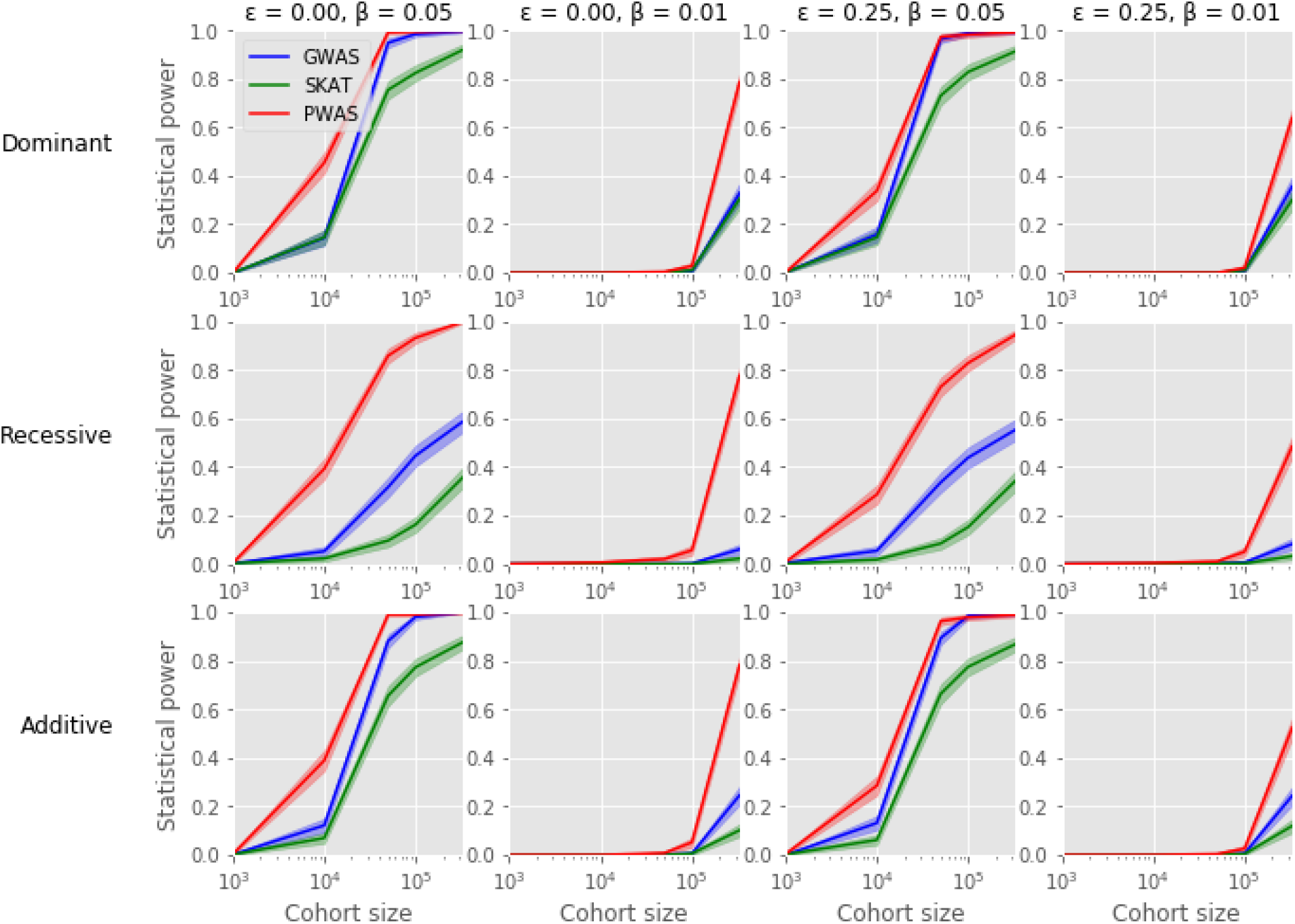
Simulation analysis. Results of a simulation analysis comparing between GWAS, SKAT and PWAS. The statistical power of each method is shown as a function of cohort size (1,000, 10,000, 50,000, 100,000 or all 332,709 filtered UKBB samples, shown in log scale). Estimated values are shown as solid lines, with flanking 95% confidence intervals as semi-transparent area bands. Each iteration of the simulation considered a single protein-coding gene affecting a simulated continuous phenotype of the form *y* = *βx* + *σ*, where *x* is the effect of the gene on the phenotype (normalized to have mean 0 and standard-deviation 1 across the UKBB population), *β* ∈ {0.01,0.05} is the gene’s effect size, and *σ*~*N*(0,1) is a random Gaussian noise. The gene effect *x* was simulated according to the PWAS model, with either a dominant, recessive or additive inheritance. A noise parameter *ϵ* ∈ {0,0.25} was introduced to FIRM, the underlying machine-learning model that estimates the damage of variants. Gene architectures, genotyping data and the 173 included covariates were taken from the UKBB cohort.

Based on the simulation results, we expect the advantage of PWAS to be the most substantial when dealing with recessive inheritance. We find that with small effect size (*β* = 0.01), at least 100K samples are required to obtain sufficient statistical power (given 173 covariates). When the effect size is higher (*β* = 0.05), cohorts of 10K samples could be sufficient.

It is important to state that phenotypes were simulated from the genetic data by a modelling scheme compatible with the assumptions of PWAS. Therefore, these results should not be seen as evidence for the dominance of PWAS over GWAS or SKAT in the real world. Rather, these simulations simply examine the method’s range of applicability and assess the amount of data required for sufficient statistical power under the settings for which it was designed. In addition to this protein-centric modeling scheme, we also examined phenotypes simulated under a standard linear model, as well as binary phenotypes (Supplementary Fig. S1).

### Case study: colorectal cancer

To examine PWAS on real phenotypes, we begin with a case-study of colorectal cancer. A cohort of 259,121 controls and 2,814 cases was derived from the UKBB to detect predisposition genes leading to increased risk of colorectal cancer through germline variants.

To exemplify how PWAS works, we begin with a demonstration of the analysis over a specific gene – *MUTYH* (Fig. 4a), a well-known predisposition gene for colorectal cancer^23^. In the studied cohort, there are 47 non-synonymous variants affecting the gene’s protein sequence. When considered by standard per-variant GWAS, the most significant of these variants yields a p-value of 1.2E-03. Even if the entire flanking region of the gene is considered (up to 500,000 bp from each side of its open reading frame), the strongest significance obtained is still only p = 6.3E-04, far from the exome-wide significance threshold (5E-07). When analyzed by PWAS, on the other hand, this association exhibits overwhelming significance (FDR q-value = 2.3E-06), far beyond the commonly used FDR significance threshold (q < 0.05).

**Fig. 4:**
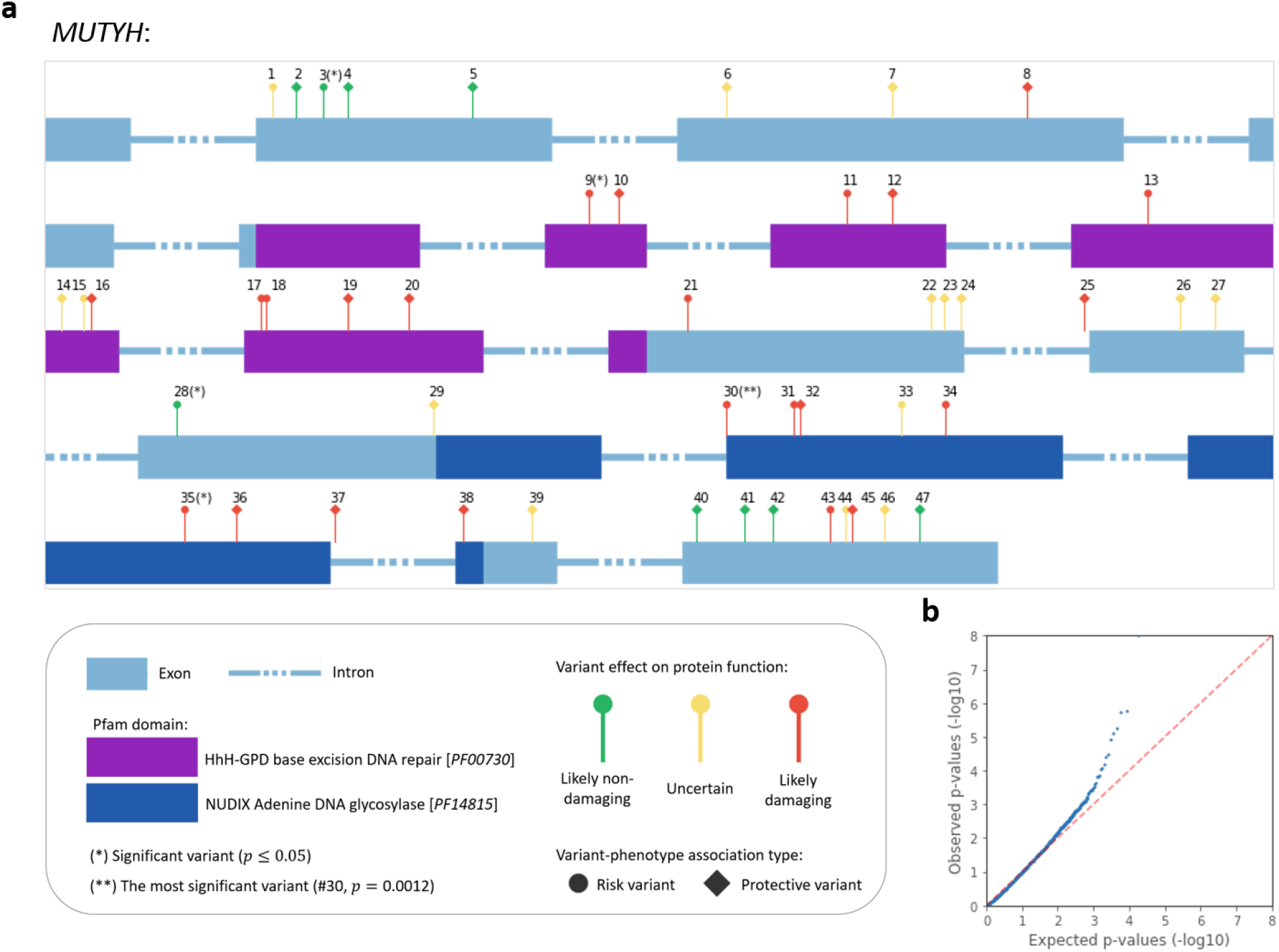
Colorectal cancer case study. (**a**) Demonstration of a specific gene-phenotype association: *MUTYH* and colorectal cancer. Variants that affect the protein sequence are shown on top of the gene’s exons. As expected, variants within domains tend to be more damaging. While none of the variants that affect the protein are close to the exome-wide significance threshold (p < 5e-07), the association is very significant by PWAS (FDR q-value = 2.3E-6). The full summary statistics of the 47 variants are presented in Supplementary Table S1. (**b**) PWAS QQ plot of all 18,053 genes tested for association with colorectal cancer.

PWAS was able to uncover the association by aggregating signal spread across a large number of different variants, with 5 of the 47 protein-affecting variants showing mild associations (p < 0.05). As these 5 variants show consistent directionality (all risk increasing), and as most of them are predicted to be likely-damaging, they were effectively aggregated into gene scores that significantly differ between cases and controls. Specifically, the *MUTYH* gene is significantly more damaged in cases than in controls according to the PWAS framework. The association is only significant according to the recessive model, with an estimated effect size of d = −0.079 (standardized mean difference in the gene effect scores between cases to controls). This observation is consistent with previous reports about *MUTYH*, claiming a recessive inheritance mode^23^.

To recover all protein-coding genes associated with colorectal cancer according to PWAS, we analyzed 18,053 genes (Fig. 4b), discovering 6 significant associations (Table 1). Of these 6 associations, 5 are supported by some literature evidence, 3 of which with level of evidence we consider strong. In 4 of the 5 supported associations, the directionality of the association reported in literature (i.e. protective or risk gene) agrees with the effect size (Cohen’s d) detected by PWAS (only in *POU5F1B* it is inversed). Of the 6 genes, only *POU5F1B* is affected by a variant exceeding the exome-wide significance (rs6998061, p = 1.4E-07). The 5 other genes are not discovered by GWAS, even when considering all the variants in the gene’s region (up to 500,000 bp away from the gene).

**Table 1:**
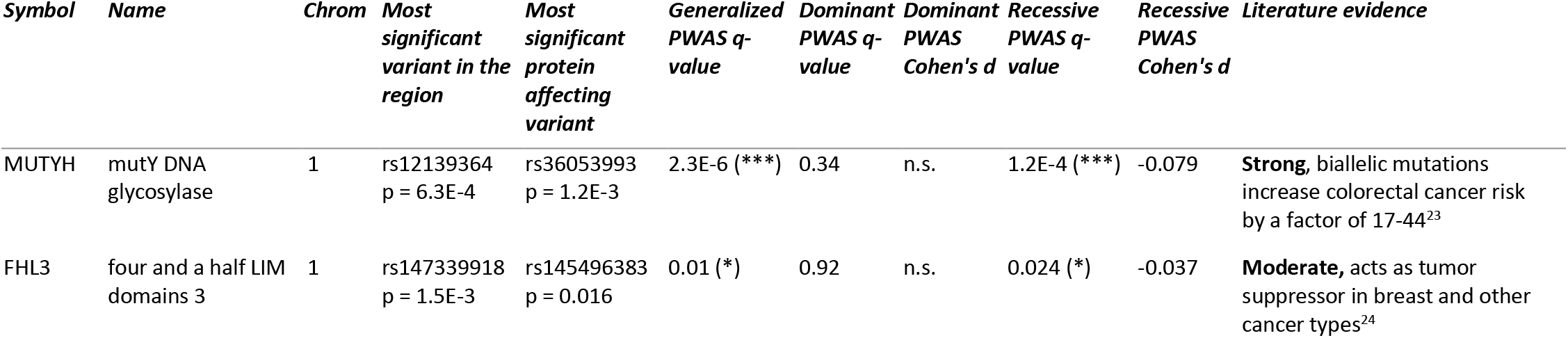
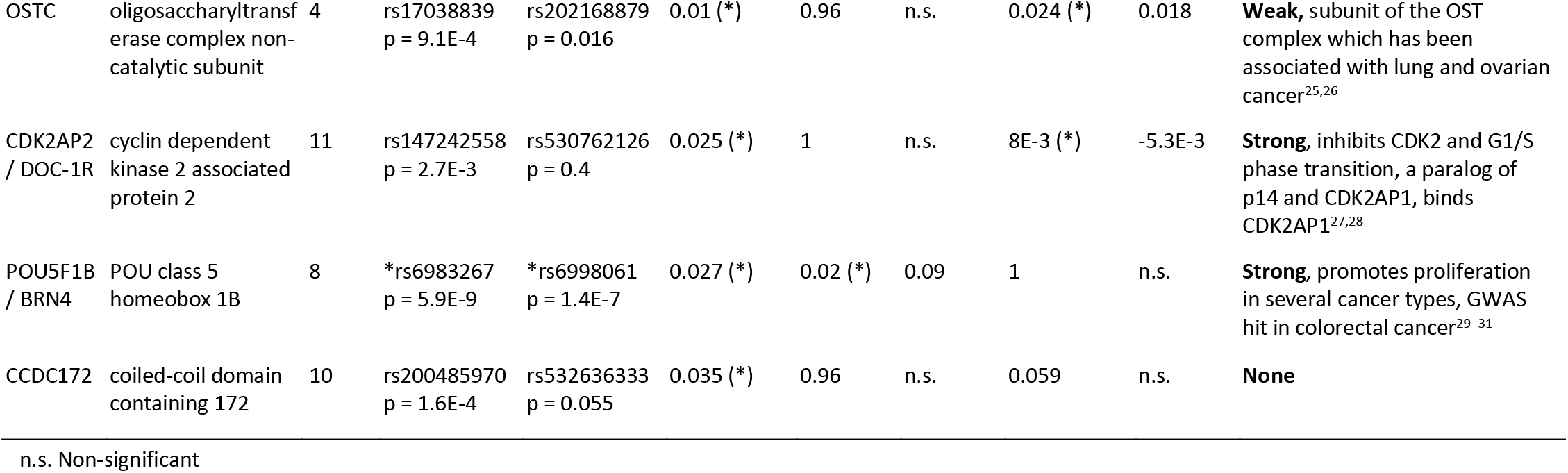
Significant colorectal cancer genes detected by PWAS

### Applicability of PWAS across 49 different phenotypes

Having case studied PWAS for a specific phenotype, we turn to consider its applicability for a diverse set of 49 prominent phenotypes (Fig. 5a). We applied both standard GWAS and PWAS across the 49 phenotypes on the same UKBB cohort (~330K samples), obtaining a rich collection of associations (Fig. 5b-c). Altogether, PWAS discovered 12,896 gene-phenotype associations, only 5,338 of which (41%) contain a GWAS-significant non-synonymous variant in the gene’s coding region (Fig. 5b). In other words, although PWAS considers the exact same set of variants, in 59% of the associations it is able to recover an aggregated signal that is overlooked by GWAS when considering each of the variants individually. Even when considering all the variants in proximity of the gene to account for LD (up to 500,000 bp to each side of the coding region), 2,998 of the 12,896 PWAS associations (23%) are still missed by GWAS (Fig. 5c-d).

**Fig. 5:**
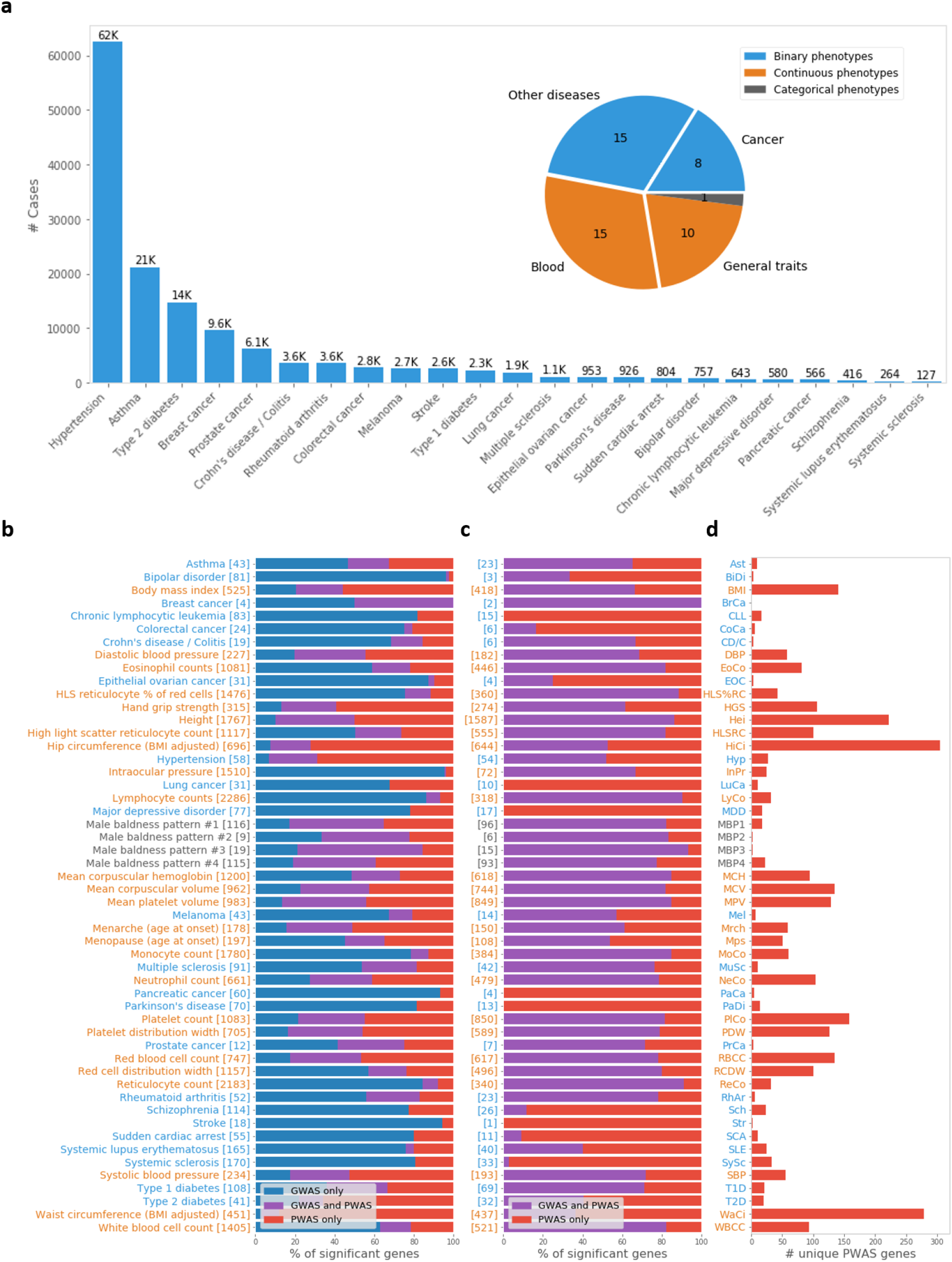
PWAS enriches GWAS discoveries across phenotypes. (**a**) We analyzed 23 binary phenotypes, 25 continuous phenotypes and 1 categorical phenotype (male-balding patterns) derived from ~330K UK Biobank samples. Within binary phenotypes, the number of cases spans orders of magnitude (from only 127 in systemic sclerosis to 62K in hypertension). (**b-c**) Partition of the significant protein-coding genes, across the different phenotypes, that were detected by GWAS, PWAS or both. The total number of significant genes is shown in brackets. In (**b**) a gene was considered significant by GWAS if a non-synonymous variant within the coding region of the gene passed the exome-wide significance threshold (p < 5E-07). In (**c**) a relaxed criterion was taken, considering all variants within 500,000 bp to each side of the coding region of the gene (here showing only the PWAS significant genes). (**d**) The number of significant genes per phenotype found by PWAS alone, according to the relaxed criterion of GWAS, as defined in (c) (i.e. without any significant variant within 500,000 bp).

Full summary of all 49 tested phenotypes, with complete per-gene summary statistics, is available in Supplementary Table S2 (for all the significant PWAS associations) and Supplementary Table S3 (with all 18,053 tested protein-coding genes). QQ plots of all 49 phenotypes are available in Supplementary Fig. S2.

To confirm the importance of the predicted functional effect scores assigned to variants, we tested the performance of a version of PWAS where the effect scores of non-synonymous variants were shuffled prior to their aggregation into gene scores. Indeed, we find that the original version of PWAS (capturing gene function) outperforms the shuffled version (Supplementary Fig. S3).

### Comparison with SKAT

Having established the discovery power of PWAS beyond standard GWAS, we also compare it to SKAT^18^, the most commonly used method for detecting genetic associations at the gene level. Importantly, whereas SKAT attempts to recover all existing genetic associations, PWAS focuses specifically on protein-coding genes that are associated with a phenotype through protein function.

We find that PWAS is superior to SKAT in the number of discovered associations for most phenotypes (Fig. 6a). We also examined the extent of overlap between the results reported by each of the two methods (the consensus associations in Fig. 6a). It appears that PWAS and SKAT tend to recover distinct sets of genes, so the two methods can be considered as largely complementary.

**Fig. 6:**
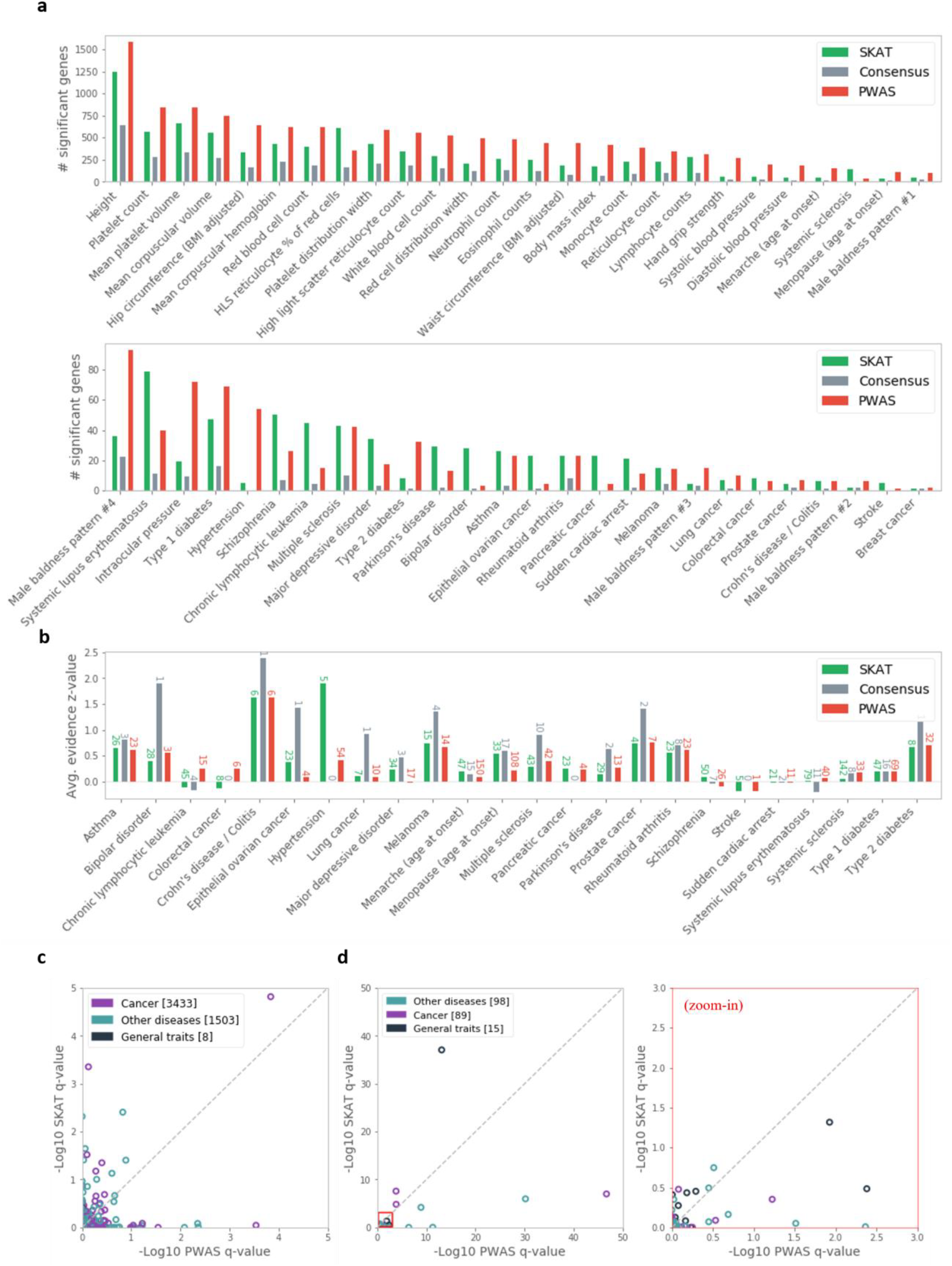
PWAS and SKAT provide complementary results. (**a**) Number of significant genes detected by PWAS, SKAT and the consensus of both, across the 49 tested phenotypes (over the same cohorts derived from the UKBB). Phenotypes are sorted by the highest of the three numbers. (**b**) An evidence score of gene-phenotype associations (derived from Open Targets Platform) is shown across phenotypes by its average over the significant genes detected by PWAS, SKAT or the consensus of both. The numbers of significant genes (over which the averaging is performed) are shown over the bars. (**c**) Comparison of the FDR q-values obtained by PWAS and SKAT over 4,944 gene-phenotype associations with strong support by Open Targets Platform. (**d**) A similar comparison over 202 associations reported by OMIM to have a known molecular basis. The right plot (marked by red frames) is a zoom-in of the left.

To assess the quality of discoveries, we appeal to Open Targets Platform (OTP)^32^, an exhaustive resource curating established gene-disease associations based on multiple layers of evidence, and OMIM^33^, the most prominent catalogue of human genes implicated in genetic disorders. We compared the quality of associations discovered by the two methods, according to OTP-derived evidence scores, across the 24 tested diseases that are recorded in OTP (Fig. 6b). According to this metric, the results of PWAS and SKAT appear to be largely comparable, with consensus genes showing stronger evidence.

We further investigate how the two methods (PWAS and SKAT) recover externally validated associations provided by OTP (Fig. 6c) and OMIM (Fig. 6d). Of 4,944 associations with strong support by OTP, 9 were recovered by SKAT compared to 6 recovered by PWAS. In the case of OMIM, which provides an even more restricted list of 202 high-quality gene-disease associations with known molecular basis, PWAS was somewhat superior (12 compared to 7 recovered associations, with the 7 being a subset of the 12). We observe no obvious trend between the types of phenotypes (e.g. cancer or other diseases) to the significance of associations obtained by the two methods (see colors in Fig. 6c-d).

Based on this comparative analysis, we conclude that PWAS and SKAT are complementary, and that it may be advantageous to use both in association studies. We stress that the two methods are very distinct in the type of associations they seek and how they model them.

### Highly-significant associations not dominated by single variants

Among all the discovered associations, we seek to highlight those that are particularly characteristic to our new method, namely results that are uniquely discovered by PWAS and show strong evidence of being causal. To this end, we filtered associations by highly strict criteria: i) strong significance (FDR q-value ≤ 0.01), ii) no other significant genes in the region, and iii) no single dominating variant association. Of the 2,998 gene-phenotype associations uniquely found by PWAS (Fig. 5c), 53 meet these criteria, and are referred to as “PWAS-exclusive” associations (Table 2; the full list is provided in Supplementary Table S4).

**Table 2:** Selected PWAS-exclusive associations

| Phenotype | Gene Symbol | Chrom | Generalized<br>q-value | Dominant<br>q-value | Dominant *r*<br>/ Cohen's *d* | Recessive<br>q-value | Recessive *r*<br>/ Cohen's *d* |
| --- | --- | --- | --- | --- | --- | --- | --- |
| Intraocular pressure | FOXG1 | 14 | 2.3E-14 | 1 | n.s. | 2.6E-15 | 0.031 |
| Hip circumference | SEMA3D | 7 | 6.3E-07 | 7.4E-05 | 0.0035 | 0.001 | 0.00034 |
| Hip circumference | ARHGAP12 | 10 | 2.1E-06 | 7.1E-07 | -0.00073 | 0.37 | n.s. |
| Colorectal cancer | MUTYH | 1 | 2.3E-06 | 0.34 | n.s. | 0.00012 | -0.079 |
| Eosinophil counts | CD80 | 3 | 3E-06 | 0.97 | n.s. | 1.1E-06 | -0.01 |
| Red cell distribution width | MLLT3 | 9 | 1.1E-05 | 0.76 | n.s. | 7.9E-06 | -0.01 |
| Hip circumference | CSGALNACT2 | 10 | 3.7E-05 | 0.00018 | -0.002 | 0.009 | -0.0021 |
| Hip circumference | C4orf36 | 4 | 4.8E-05 | 2.3E-05 | -0.0019 | 0.00014 | 0.0038 |
| Type 2 diabetes | POU3F4 | X | 0.0016 | 0.00052 | 0.04 | N/A | N/A |
| Type 2 diabetes | USP26 | X | 0.0016 | 0.015 | 0.049 | N/A | N/A |
| Chronic lymphocytic leukemia | FAM160B1 | 10 | 0.0033 | 0.99 | n.s. | 0.048 | 0.06 |

n.s. Non-significant; N/A not applicable (X-linked recessive inheritance)

As expected, the PWAS-exclusive genes show no GWAS signal at all, and the PWAS associations are constrained to the associated genes (Fig. 7a). When considered by SKAT, only 3 of the 53 associations come up as significant (Fig. 7b), even though SKAT was not included in the criteria for defining those associations.

**Fig. 7:**
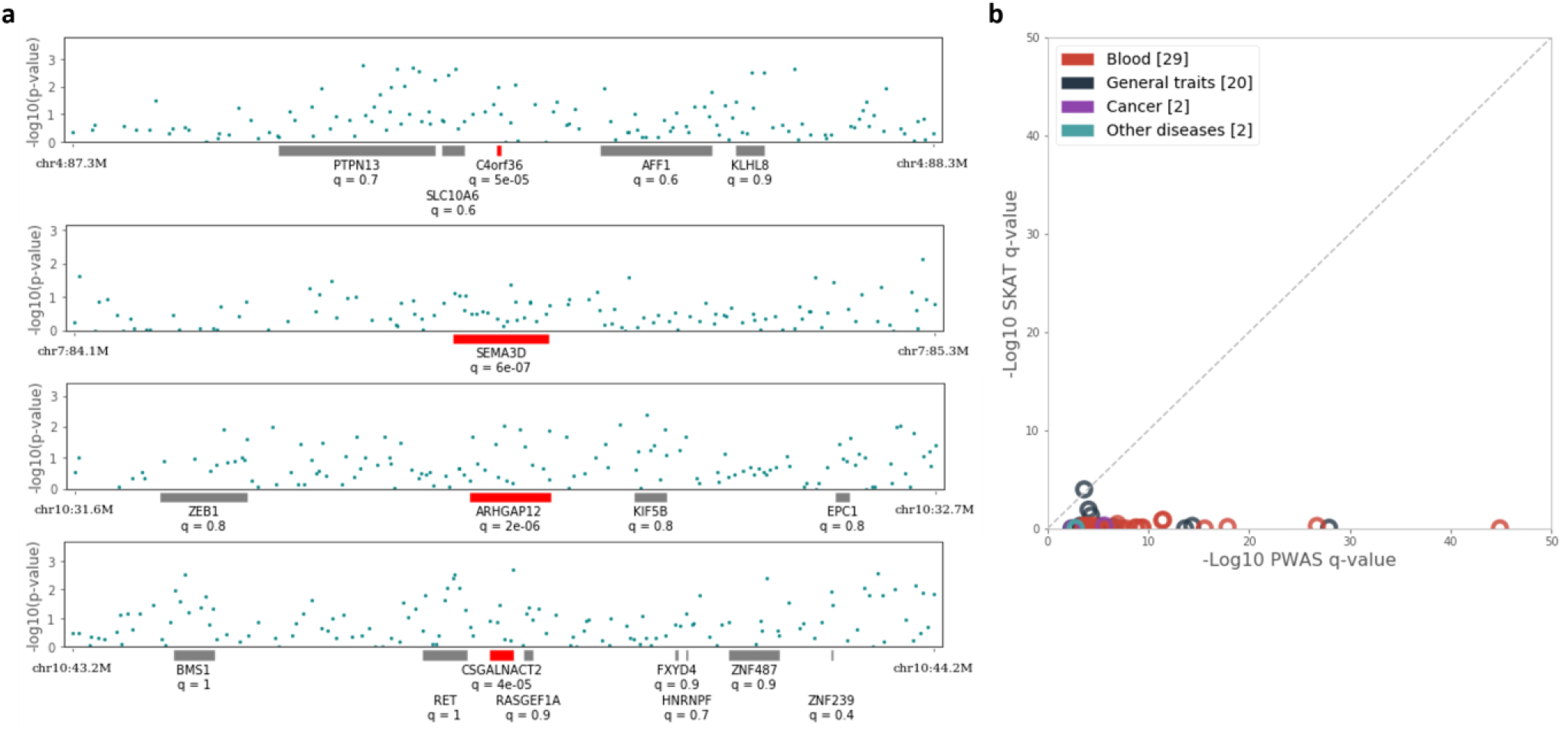
PWAS-exclusive associations. (**a**) Exemplifying the 53 PWAS-exclusive associations with the 4 genes associated with the hip circumference phenotype. The 4 genes demonstrate a complete lack of any GWAS pattern in proximity of the genes (up to 500,000 bp to both directions of each gene). Each of the 4 depicted gene regions was divided into 200 bins, displaying the most significant variant in each bin. Also shown are the PWAS FDR q-values of all analyzed protein-coding genes in those chromosomal regions. (**b**) Comparison of the FDR q-values obtained by PWAS and SKAT for the 53 associations.

Many of the listed associations are strongly support by literature. For example, *POU3F4* (also known as *BRN4*) was found by PWAS to be associated with type 2 diabetes (FDR q-value = 0.0016), apparently as a protective gene (Cohen’s d of 0.04 and 0.033 according to the dominant and recessive models, respectively). *BRN4* is an essential gene for the development of pancreatic α-cells, whose excess glucagon secretion is implicated in type-2 diabetes^34^. In other words, impairment of the gene is expected to reduce glucagon levels, making it protective of the disease.

MLLT3, appearing to be associated with red blood cell distribution width through recessive inheritance according to PWAS (FDR q-value = 7.9E-06, r = −0.01), has been reported to be a key regulatory gene in the bone marrow^35^. Likewise, *CD80*, which PWAS associates with eosinophil counts through recessive inheritance (FDR q-value = 1.1E-06, r = −0.01), has an important role in antigen presentation by eosinophils^36^.

In other cases, while there is no clear indication for the reported association, there does exist a strong molecular plausibility. *FOXG1*, for example, plays a key role in the development of the retina (a function conserved in all vertebrates)^37^, and was shown to be associated with visual impairment in both mice and human cohorts^38^. However, it has never been implicated in intraocular pressure, an association that we observe here with extraordinary significance (FDR q-value = 2.6E-15) according to the PWAS recessive model. Specifically, normal function of the gene (i.e. lack of damaging variants) appears to be positively correlated (r = 0.031) with intraocular pressure.

In some instances, we find little to no literature evidence for reported PWAS-exclusive associations (e.g. *C4orf36* and Hip circumference, *FAM160B1* and leukemia, *USP26* and type 2 diabetes), yet the strong associations established by PWAS provide strong evidence for these associations.

## Discussion

In this work, we have introduced a new functional protein-centric approach to association studies. We have demonstrated its applicability to a broad range of prominent human phenotypes, and established its utility in supplementing existing methods and highlighting novel associations.

Due to its explicit gene-based functional model, PWAS provides more interpretable results than other methods. Like other gene-based approaches seeking to establish associations of concrete genes, it requires no post-analysis fine-mapping. Furthermore, as PWAS relies on an explicit functional model, it is better posed to suggest causal relationships. Specifically, a significant PWAS association would suggest that variants disrupting the function of the implicated protein might influence the studied phenotype (in the case of a disease, increase or decrease one’s risk). Furthermore, PWAS can determine whether the proposed causal effect appears to be dominant, recessive or some mixture of the two (e.g. additive). Yet, while PWAS is more suggestive of causality than other methods, significant results are still susceptible to spurious correlations. In particular, the problem of LD^7^ is still far from being resolved, and significant PWAS associations, like any genetic associations, should be interpreted with caution.

By aggregating all variants affecting the same gene into unified statistics, PWAS is able to detect signal that is too weak and spread to appear in per-variant GWAS (Fig. 7, Table 2). It is particularly important in the case of rare variants, which account for much of the heritability^39^. In fact, PWAS can successfully handle even variants that occur only once in the cohort (including, in principle, de-novo variants). As long as the observed variants fit the overall trend observed in the studied gene (e.g. that they are more damaging in cases compared to controls), even singletons can increase the statistical power of the method. In this work, however, we relied on imputed genotypes which cannot capture variants that are too rare. As a result, some biological signals have probably been missed (e.g. damaged genes that were mistaken to be intact due to ungenotyped variants). We therefore anticipate that PWAS can substantially benefit from exome sequencing (as opposed to SNP-array genotypes). It should be noted that while more accurate genotyping should indeed enhance its statistical power, the reliance of PWAS on rigorous statistics keeps it insensitive to false discoveries even with imperfect genotyping.

A rather unique feature of PWAS is its separate dominant and recessive inheritance models. Although there are strong indications that the commonly used additive model can capture most of the heritability of complex human traits^40^, non-additive and epistatic effects play key role in many phenotypes^41^. While there have been efforts to address epistatic effects in GWAS^42^, the special case of recessive inheritance in complex traits has been largely neglected. Our results show that recessive inheritance is indeed substantial in a variety of phenotypes. 23% of the recovered PWAS associations are significant by the recessive but not the dominant model. PWAS is uniquely posed, among present methods, to handle recessive inheritance, as per-gene recessive inheritance is much more sensible than per-variant. Specifically, PWAS is able to capture the prevalent instances of compound heterozygosity (due to its per-gene aggregation), whereas per-variant GWAS would fail to detect such recessive effects^43^.

Another important advantage of PWAS over existing methods is its reduced computational burden in multi-phenotype datasets (such as the UKBB). PWAS aggregates all the genetic data into compact gene score matrices, whose size is much smaller than the raw genotyping data (as there are typically substantially fewer genes than variants). These matrices store all of the relevant genetic information (encompassing the assessed functional state of the proteome in each of the cohort samples), and can be independently tested against each phenotype.

PWAS belongs to the growing family of methods that seek genetic associations through modeling of functional genomic properties. While methods such as PrediXcan^20^ and TWAS^21^ model gene expression, PWAS models protein function, which, in principle, is completely orthogonal to the signal of gene abundance. We purposefully employ a very abstract definition of the term “protein function” to encapsulate anything the protein is supposed to do in the cell such that disturbing it (by variants altering the protein sequence) could lead to phenotypic effects (e.g. missense variants affecting a membrane receptor protein could interfere with its signal transduction function and result in predisposition to cancer). We consider PWAS complementary to methods that model other functional aspects of the genome.

Contrary to expression-based methods, PWAS assigns protein effect scores in a deterministic, consistent manner. Gene expression is highly volatile, with substantial variability between tissues, epigenetic conditions and many other non-genetic factors. In contrast, protein products are mostly a direct result of one’s genetic makeup. This benefits PWAS in two major ways. First, it offers reduced computational complexity, since it is sufficient to compute the gene score matrices only once. More importantly, it relieves us from the need to select a specific tissue or expression profile for the analysis. Indeed, most human traits are not confined to specific tissues, let alone specific cellular conditions, making the selection of a relevant reference panel for expression-based methods a daunting task.

A potential limitation of PWAS is its reliance on the complete cohort data (including the full genotype and phenotype information). Unlike other methods, it is unable to analyze summary GWAS statistics alone. This reliance on raw data is due to the non-linear nature of the aggregation algorithm used to derive gene effect scores from variant effect scores (see Methods). It remains open whether a simplified linear version of PWAS could be derived, or at least a version simple enough that can be applied on summary statistics. On the positive side, with modern biobanks and genetic cohorts (e.g. UKBB, SFARI^44^), large-scale datasets are becoming increasingly available for direct modeling and analysis.

In conclusion, we have presented PWAS as a novel protein-centric method for genetic association studies providing functionally interpretable gene results. We have demonstrated the validity of PWAS through comparison to multiple external resources, and shown its added value to commonly used methods across a wide range of prominent phenotypes, including numerous new discoveries. We argue that integrating rich machine-learning models based on prior-knowledge, as exemplified in this work, is a promising avenue to novel insight and discovery in human genetics.

## Methods

### UK Biobank cohort

Throughout this work we used genetic and phenotypic data from the UK Biobank (UKBB) resource^8,9^ (application ID 26664).

From the entire UKBB cohort of 502,539 samples, we filtered 409,600 labeled as Whites/Caucasians according to both self-reported ethnicity and their genetics. We removed 312 samples with mismatching self-reported and genetics-derived sex. We also removed 726 samples without imputed genotypes.

Finally, we removed 75,853 samples to keep only one representative of each kinship group of related individuals, obtaining a final cohort of 332,709 samples.

Specification of the 49 phenotypes used in this work is available in Supplementary Table S5. The table specifies how each phenotype was defined (based on either a UKBB field or ICD-10 codes), and whether it was restricted to a specific gender. The set of all ICD-10 codes associated with a sample were derived from the following UKBB fields: 41202, 41204, 40006, 40001, 40002, 41201.

When testing a specific phenotype, we also filtered out samples with missing values in that phenotype (e.g. for height we filtered out 686 samples, obtaining a cohort of 332,023 samples). When testing phenotypes defined by ICD-10 codes, we filtered out all samples without any recorded ICD-10 code. This further removed 70,335 samples from the cohort, leaving 262,374 samples in those phenotypes. The final cohort size used for testing each phenotype is listed in Supplementary Table S2. In the rare cases where samples had multiple records of the same continuous phenotype (e.g. from different visits to the UKBB assessment centers), we took the maximum value.

All the association tests carried out in this work (with either of the three used methods, i.e. GWAS, PWAS or SKAT) included the following covariates: sex (binary), year of birth (numeric), the 40 principal components of the genetic data provided by the UKBB (numeric), the UKBB genotyping batch (one-hot-encoding with 105 categories) and the UKBB assessment centers associated with each sample (binary, with 25 categories). Altogether, 173 covariates (including a constant intercept) were included. For specific phenotypes, additional covariates were included as part of the phenotype’s definition (e.g. “Hip circumference adjusted for BMI” included BMI as an additional covariate; see Supplementary Table S5).

### Variant functional effect scores

The gene effect scores used by PWAS are derived from aggregation of per-variant effect scores (Fig. 2b). Each non-synonymous variant in the coding region of a gene which affects the resulted protein sequence is assigned a functional effect score that aims to capture its propensity to damage the protein product of the gene. Specifically, PWAS considers the following types of variants as affecting protein sequence: missense, nonsense, frameshift, in-frame indel and canonical splice-site variants. The predicted effect score of a variant is a number between 0 (complete loss of function) to 1 (no functional effect). Intuitively, it reflects the probability that the affected gene retains its function given the variant.

To predict the effect of missense variants, PWAS employs a machine-learning model. Specifically, the FIRM predictor is used^22^. Unlike commonly used prediction tools assessing mutation pathogenicity (e.g. CADD^45^, SIFT^46^, Polyphen2^47^, MutationTaster2^48^), FIRM is designed to assess the damage of variants at the molecular level (rather than clinical outcome at the organism level). This distinction is particularly important when PWAS is used for phenotypes without clinical significance (e.g. height). FIRM attempts to capture gene function in its broadest sense (e.g. enzymatic reaction, molecular interaction, cellular pathways), thereby allowing PWAS, in principle, to model any protein-phenotype effect.

To assess the impact of a missense variant on gene function, FIRM considers its rich proteomic context, which it encodes as a set of 1,109 numerical features (which are used by the underlying machine-learning model to predict its effect score). The full specification of the features used by FIRM is described elsewhere^22^. They include: i) the position of the variant within the protein sequence, ii) the identity of the reference and alternative amino-acids and the amino-acid composition of the protein in various regions of the protein with respect to the variant, iii) abundance of annotations extracted from UniProt^49^ (e.g. phosphorylation and other post-translational modifications, active sites, secondary structure), and iv) details of Pfam domains^50^ in proximity of the variant.

Missense variants comprise the vast majority of non-synonymous variants^51^. For other variant types, effect scores were derived through rougher, rule-based formulas. Specifically, nonsense, frameshift and canonical splice-site variants (i.e. variants affecting the first or last two letters of an intron) were assumed to be loss-of-function variants and assigned a score of 0. In-frame indels were assigned an effect score based on the numbers of substituted, inserted and deleted amino-acids (see Supplementary Methods).

### Gene functional effect scores

To calculate gene effect scores (Fig. 1b, Fig. 2c-d), PWAS aggregates variant effect scores (see previous section) integrated with the genotyping data. Unlike variant-level scores, gene scores are sample specific (i.e. depending on each sample’s genotype). PWAS supports two aggregation schemes, resulting in “dominant” and “recessive” gene scores. Intuitively, dominant scores reflect the probability of at least one damaging event, whereas recessive scores reflect the probability for at least two.

Let *s*_1_, …, *s*_*k*_ be the functional effect scores assigned to the *k* variants potentially affecting a protein-coding gene by the scheme detailed in the previous section. For a given variant *i* ∈ [*k*] (in the context of a given sample), let 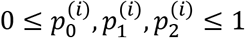 (satisfying 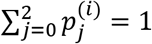) indicate the genotyping probabilities of the variant (i.e. 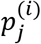 is the probability of variant *i* to occur *j* times in the given sample). Recall that we intuitively interpret *s*_*i*_ as the probability that the gene retains its function following the variant effect.

A question arises how to estimate the probability of the gene to retain its functions if variant *i* occurs twice (an event of probability 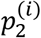). A possible approach would be to treat the two occurrences of the variant as independent, so the probability would be 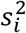. Another approach is to treat the two occurrences as fully dependent (i.e. either the variant is damaging or it isn’t), taking the probability to be simply *s*_*i*_ like in the heterozygous case. To accommodate this uncertainty, we chose to introduce a parameter *μ* ∈ [0,1] and take the effect to be 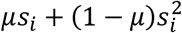. The parameter *μ* can be thought of as the probability of the homozygous effect to be dependent (i.e. when *μ* = 0 it is completely independent, and when *μ* = 1 it is fully dependent). Overall, the probability that the gene retains its function considering variant *i* (in the context of that sample) would be 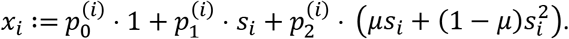.

Note that in reality the scores *s*_*i*_ are not purely probabilistic entities. More likely, they capture both the probability of gene damage and its extent (so *s*_*i*_ and *x*_*i*_ can be more realistically interpreted as damage expectations rather than probabilities). That is another reason why the independent case (*μ* = 0) might be more appropriate than the dependent case (*μ* = 1), as two hits of a variant often cause more damage than a single hit. Taking the same expression (*s*_*i*_) to estimate the outcome of these two events would miss this effect.

We want the dominant effect score of the gene to reflect the probability that it retains its function (given the sample’s genotyping of the *k* variants and their effect scores). If we simplistically assume that the *k* variants independently affect the gene, then it retains its function with probability *x*_1_ … *x*_*k*_. Here too, some degree of dependence might better reflect the dominant effect of the gene. Under full dependence, we would take the score to be min{*x*_1_, …, *x*_*k*_} (i.e. the overall effect on the gene is the effect of the most damaging variant). To allow a more refined dependence model, let us write the multiplication 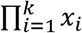 as 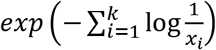. The term 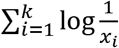 is the ℓ_1_ norm of the vector 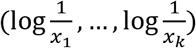. We introduce another parameter 1 ≤ *p* ≤ ∞ and take the dominant score to be 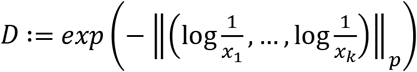. Note that when *p* = ∞ we get the full independence score min{*x*_1_, …, *x*_*k*_}.

For deriving the recessive effect score of the gene, we would like to express the probability of at most one damaging event (so its complementary event would represent the probability of at least two damaging events). Assuming independence, that probability would be 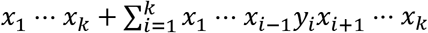, where *y*_*i*_ expresses the probability of variant *i* damaging exactly one copy of the gene. Specifically, we define 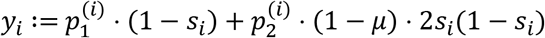. The second coefficient is explained by 2*s*_*i*_(1 − *s*_*i*_) being the probability of variant *i* introducing exactly one hit, given that each of its two copies are independent; when they are fully dependent, that is not possible for the two copies to introduce exactly one hit. When all *x*_*i*_ ≠ 0, we can rewrite that expression as 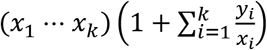. Like with the dominant score, we parameterize (*x*_*i*_ … *x*_*k*_) into 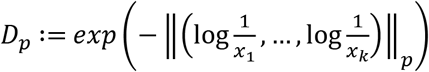, and 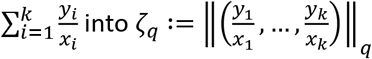 for some parameter values 1 ≤ *p*, *q* ≤ ∞. The recessive effect score is then taken to be (1 + ζ_*q*_)*D*_*p*_. However, this term is not well-defined when there is *x*_*i*_ = 0. To derive the recessive score in that case, we can calculate 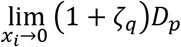 (see Supplementary Methods for details) and obtain:

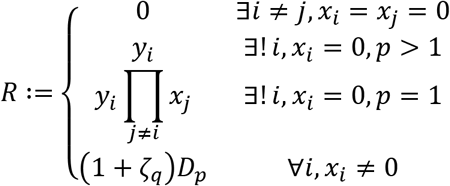

To summarize, the aggregation scheme takes as input the individual variant scores *s*_1_, …, *s*_*k*_ (which are sample independent) and the genotyping probabilities of the *k* variants within the given sample 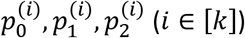, to produce the dominant and recessive gene scores of the gene. The dominant score *D* relies on two parameters (*μ* and *p*), whereas the recessive score *R* depends on three parameters (*μ*, *p* and *q*). Note that the parameters *μ* and *p* used by the two scoring schemes need not take the same values in the two contexts (despite sharing a similar purpose). For clarity, we denote the parameters of *D* by *μ*_*D*_ and *p*_*D*_, and the parameters of *R* by *μ*_*R*_, *p*_*R*_ and *q*_*R*_. Overall, the effect score aggregation scheme of PWAS is parameterized by 5 distinct parameters.

To find optimal parameter values, we fit the aggregation scheme on known gene-phenotype associations derived from OMIM^33^, taking the combination of 5 parameters that optimize the recovered significance of these associations (see Supplementary Methods). Importantly, the gene-phenotype associations used to find the optimal parameters do not overlap with the associations used to evaluate PWAS throughout this work (e.g. Fig. 6c). In particular, they involve other phenotypes that were not studied in the primary analysis. The obtained parameter values used throughout the analyses presented in this work are: *μ*_*D*_ = 1, *p*_*D*_ = 1.25, *μ*_*R*_ = 0.5, *p*_*R*_ = ∞, *q*_*R*_ = 3.

### Non-modeled genomic properties

In its current form, PWAS does not consider structural and copy number variations, as they do not naturally fit into the framework of dominant and recessive heritability modes. Non-canonical splicing effects are also not considered at present, as they are not supported by FIRM. In general, the effects of splicing events are considered to be hard to model^52^. Furthermore, weak splicing events are often associated with alternative splicing of non-canonical protein isoforms. To allow simple modeling and interpretation of the results, PWAS considers only canonical protein isoforms (see Supplementary Methods).

It should also be noted that the recessive model assumes standard autosomal inheritance, and PWAS does not properly address recessive inheritance in sex and mitochondrial chromosomes. Another current limitation of the recessive model has to do with the absence of phased genotypes in the UKBB resource. For a recessive genetic effect to take place, both copies of a gene (on the two copies of the relevant chromosome) should be affected. Due to the lack of phased genotypes, PWAS is unable to determine if different variants affect the same or different copies of the gene. Therefore, our modeling choice was to assume that different variants affect different gene copies (see previous section).

Importantly, these non-modeled genomic properties can only affect the statistical power of PWAS, but should not lead to false discoveries (see next section).

### Statistical analysis

To test whether a gene is associated with a phenotype, PWAS conducts linear or logistic regression (depending on whether the phenotype is continuous or binary, respectively). A categorical phenotype is split into multiple binary phenotypes (each isolating one of the categories in a one-vs.-rest manner). The regression model includes all 173 covariates (see the “UK Biobank dataset” section), and the relevant gene scores (dominant, recessive, or both). Specifically, when testing for dominant inheritance, the term *β*_*D*_ ⋅ *D* is included in the regression model, where *D* is the dominant score of the gene, and *β*_*D*_ is the corresponding regression coefficient. The null hypothesis of the regression under dominant inheritance is *H*_0_: *β*_*D*_ = 0. Similarly, when testing for recessive inheritance, the term *β*_*R*_ ⋅ *R* is included, and the null hypothesis is *H*_0_: *β*_*R*_ = 0. When the test is carried according to the generalized model, both terms are included in the regression, and the tested null hypothesis is *H*_0_: *β*_*D*_ = *β*_*R*_ = 0. Unless stated explicitly that the dominant or recessive model is used, all the p-values reported in this work refer to the generalized model.

As PWAS relies on routine statistical analysis to calculate significance, its results are valid (in terms of avoiding false discoveries) regardless of how accurately the calculated gene scores reflect the true underlying biology. While better scoring schemes are expected to provide increased statistical power, protection against type-I errors is guaranteed irrespectively.

To provide a fair comparison to PWAS, the results of GWAS and SKAT reported in this work were performed using identical statistical analysis over the same data (see Supplementary Methods).

### Source code availability

An effort is currently underway to organize and document the source code of PWAS and provide it as an open-source project in GitHub with command-line interface. We expect this effort to be completed shortly. Meanwhile, the source code is available upon request.

## Supplementary materials

**Supplementary Table S1**: The 47 imputed variants affecting the protein sequence of the *MUTYH* gene.

**Supplementary Table S2**: Full per-gene summary statistics of all significant PWAS associations across all 49 tested phenotypes.

**Supplementary Table S3**: Full per-gene summary statistics of all 18,053 tested protein-coding genes across all 49 tested phenotypes.

**Supplementary Table S4**: The 53 gene-phenotype associations defined as “PWAS-exclusive”.

**Supplementary Table S5**: Specification of the 49 tested phenotypes.

**Supplementary Table S6**: Specification of the 6 OMIM diseases used to fit the effect score aggregation models.

**Supplementary Table S7**: Enrichment factor between our GWAS results to the GWAS Catalog across all 49 tested phenotypes and parameter values.

**Supplementary Fig. S1:** Simulation analysis under binary and linear models.

**Supplementary Fig. S2:** PWAS QQ plots of all phenotypes.

**Supplementary Fig. S3:** PWAS with shuffled effect scores.

**Supplementary Fig. S4:** Distribution of combined p-values over the explored parameters of the effect score aggregation models.

**Supplementary Fig. S5:** QQ plot of the explored parameters of the effect score aggregation models.

**Supplementary Fig. S6:** GWAS Manhattan plots of selected phenotypes.

**Supplementary Fig. S7:** GWAS QQ plots of all phenotypes.

**Supplementary methods**

## References

1. Lewis, C. M. & Knight, J. Introduction to genetic association studies. Cold Spring Harb. Protoc. 2012, pdb--top068163 (2012).

2. Korte, A. & Farlow, A. The advantages and limitations of trait analysis with GWAS: a review. Plant Methods 9, 29 (2013).

3. Loh, P.-R. et al. Efficient Bayesian mixed-model analysis increases association power in large cohorts. Nat. Genet. 47, 284 (2015).

4. Chang, C. C. et al. Second-generation PLINK: rising to the challenge of larger and richer datasets. Gigascience 4, 7 (2015).

5. Buniello, A. et al. The NHGRI-EBI GWAS Catalog of published genome-wide association studies, targeted arrays and summary statistics 2019. Nucleic Acids Res. 47, D1005–D1012 (2018).

6. Visscher, P. M., Brown, M. A., McCarthy, M. I. & Yang, J. Five years of GWAS discovery. Am. J. Hum. Genet. 90, 7–24 (2012).

7. Visscher, P. M. et al. 10 years of GWAS discovery: biology, function, and translation. Am. J. Hum. Genet. 101, 5–22 (2017).

8. Sudlow, C. et al. UK biobank: an open access resource for identifying the causes of a wide range of complex diseases of middle and old age. PLoS Med. 12, e1001779 (2015).

9. Bycroft, C. et al. Genome-wide genetic data on ~500,000 UK Biobank participants. BioRxiv 166298 (2017).

10. Manolio, T. A. et al. Finding the missing heritability of complex diseases. Nature 461, 747 (2009).

11. Spain, S. L. & Barrett, J. C. Strategies for fine-mapping complex traits. Hum. Mol. Genet. 24, R111–R119 (2015).

12. Schaid, D. J., Chen, W. & Larson, N. B. From genome-wide associations to candidate causal variants by statistical fine-mapping. Nat. Rev. Genet. 19, 491 (2018).

13. de Bunt, M. et al. Evaluating the performance of fine-mapping strategies at common variant GWAS loci. PLoS Genet. 11, e1005535 (2015).

14. Zhu, Z. et al. Integration of summary data from GWAS and eQTL studies predicts complex trait gene targets. Nat. Genet. 48, 481 (2016).

15. Hormozdiari, F. et al. Colocalization of GWAS and eQTL signals detects target genes. Am. J. Hum. Genet. 99, 1245–1260 (2016).

16. Pers, T. H. et al. Biological interpretation of genome-wide association studies using predicted gene functions. Nat. Commun. 6, 5890 (2015).

17. Hormozdiari, F., Kichaev, G., Yang, W.-Y., Pasaniuc, B. & Eskin, E. Identification of causal genes for complex traits. Bioinformatics 31, i206–i213 (2015).

18. Wu, M. C. et al. Rare-variant association testing for sequencing data with the sequence kernel association test. Am. J. Hum. Genet. 89, 82–93 (2011).

19. Lee, S. et al. Optimal unified approach for rare-variant association testing with application to small-sample case-control whole-exome sequencing studies. Am. J. Hum. Genet. 91, 224–237 (2012).

20. Gamazon, E. R. et al. A gene-based association method for mapping traits using reference transcriptome data. Nat. Genet. 47, 1091 (2015).

21. Gusev, A. et al. Integrative approaches for large-scale transcriptome-wide association studies. Nat. Genet. 48, 245 (2016).

22. Brandes, N., Linial, N. & Linial, M. Quantifying gene selection in cancer through protein functional alteration bias. Nucleic Acids Res. (2019).

23. Lubbe, S. J., Di Bernardo, M. C., Chandler, I. P., Houlston, R. S. & others. Clinical implications of the colorectal cancer risk associated with MUTYH mutation. J Clin Oncol 27, 3975–3980 (2009).

24. Niu, C. et al. Downregulation and antiproliferative role of FHL3 in breast cancer. IUBMB Life 63, 764–771 (2011).

25. Sambrooks, C. L. et al. Oligosaccharyltransferase Inhibition Overcomes Therapeutic Resistance to EGFR Tyrosine Kinase Inhibitors. Cancer Res. 78, 5094–5106 (2018).

26. Va\vnhara, P. et al. Loss of the oligosaccharyl transferase subunit TUSC3 promotes proliferation and migration of ovarian cancer cells. Int. J. Oncol. 42, 1383–1389 (2013).

27. Liu, Q. et al. Overexpression of DOC-1R inhibits cell cycle G1/S transition by repressing CDK2 expression and activation. Int. J. Biol. Sci. 9, 541 (2013).

28. Ertekin, A. et al. Human cyclin-dependent kinase 2-associated protein 1 (CDK2AP1) is dimeric in its disulfide-reduced state, with natively disordered N-terminal region. J. Biol. Chem. 287, 16541–16549 (2012).

29. Hayashi, H. et al. The OCT4 pseudogene POU5F1B is amplified and promotes an aggressive phenotype in gastric cancer. Oncogene 34, 199 (2015).

30. Pan, Y. et al. POU5F1B promotes hepatocellular carcinoma proliferation by activating AKT. Biomed. Pharmacother. 100, 374–380 (2018).

31. Yao, L., Tak, Y. G., Berman, B. P. & Farnham, P. J. Functional annotation of colon cancer risk SNPs. Nat. Commun. 5, 5114 (2014).

32. Carvalho-Silva, D. et al. Open Targets Platform: new developments and updates two years on. Nucleic Acids Res. 47, D1056–D1065 (2018).

33. Hamosh, A., Scott, A. F., Amberger, J. S., Bocchini, C. A. & McKusick, V. A. Online Mendelian Inheritance in Man (OMIM), a knowledgebase of human genes and genetic disorders. Nucleic Acids Res. 33, D514–D517 (2005).

34. Phadnis, S. M., Ghaskadbi, S. M., Hardikar, A. A. & Bhonde, R. R. Mesenchymal stem cells derived from bone marrow of diabetic patients portrait unique markers influenced by the diabetic microenvironment. Rev. Diabet. Stud. RDS 6, 260 (2009).

35. Pina, C., May, G., Soneji, S., Hong, D. & Enver, T. MLLT3 regulates early human erythroid and megakaryocytic cell fate. Cell Stem Cell 2, 264–273 (2008).

36. Tamura, N. et al. Requirement of CD80 and CD86 molecules for antigen presentation by eosinophils. Scand. J. Immunol. 44, 229–238 (1996).

37. Picker, A. et al. Dynamic coupling of pattern formation and morphogenesis in the developing vertebrate retina. PLoS Biol. 7, e1000214 (2009).

38. Boggio, E. M. et al. Visual impairment in FOXG1-mutated individuals and mice. Neuroscience 324, 496–508 (2016).

39. Speed, D. et al. Reevaluation of SNP heritability in complex human traits. Nat. Genet. 49, 986 (2017).

40. Hill, W. G., Goddard, M. E. & Visscher, P. M. Data and theory point to mainly additive genetic variance for complex traits. PLoS Genet. 4, e1000008 (2008).

41. Moore, J. H. & Williams, S. M. Epistasis and its implications for personal genetics. Am. J. Hum. Genet. 85, 309–320 (2009).

42. Niel, C., Sinoquet, C., Dina, C. & Rocheleau, G. A survey about methods dedicated to epistasis detection. Front. Genet. 6, 285 (2015).

43. Gibson, G. Rare and common variants: twenty arguments. Nat. Rev. Genet. 13, 135 (2012).

44. Feliciano, P. et al. SPARK: a US cohort of 50,000 families to accelerate autism research. Neuron 97, 488–493 (2018).

45. Kircher, M. et al. A general framework for estimating the relative pathogenicity of human genetic variants. Nat. Genet. 46, 310 (2014).

46. Ng, P. C. SIFT: predicting amino acid changes that affect protein function. Nucleic Acids Res. 31, 3812–3814 (2003).

47. Adzhubei, I., Jordan, D. M. & Sunyaev, S. R. Predicting functional effect of human missense mutations using PolyPhen-2. Curr. Protoc. Hum. Genet. 76, 7–20 (2013).

48. Schwarz, J. M., Cooper, D. N., Schuelke, M. & Seelow, D. MutationTaster2: mutation prediction for the deep-sequencing age. Nat. Methods 11, 361 (2014).

49. Consortium, U. UniProt: a hub for protein information. Nucleic Acids Res. 43, D204–D212 (2014).

50. Finn, R. D. et al. Pfam: the protein families database. Nucleic Acids Res. 42, D222–30 (2014).

51. Karczewski, K. J. et al. Variation across 141,456 human exomes and genomes reveals the spectrum of loss-of-function intolerance across human protein-coding genes. BioRxiv 531210 (2019).

52. Stamm, S. et al. Function of alternative splicing. Gene 344, 1–20 (2005).

