## Supplementary Materials for "PWAS: Proteome-Wide Association Study"

### Supplementary figures

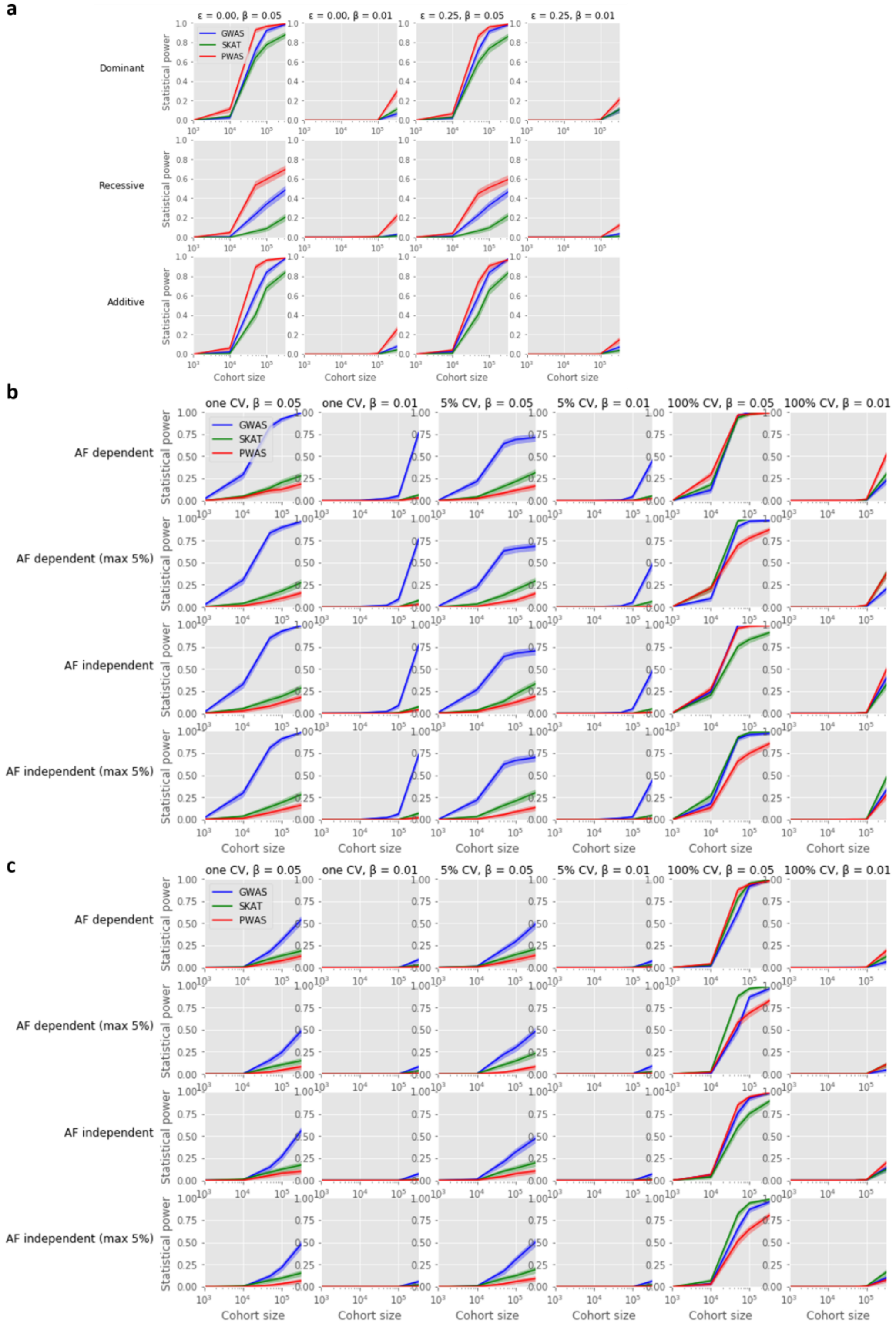

**Supplementary Fig. S1: Simulation analysis under binary and linear models**

(**a**) An extension of the protein-centric modeling scheme (Fig. 3) to binary phenotypes. For simulating binary phenotypes, the continuous result $\beta x + \sigma$ is binarized into $y = \theta(\beta x + \sigma)$, where $\theta$ is a step function. (**b**) In addition to the protein-centric simulation model (which suits the assumptions of PWAS), we also explored a standard linear model conforming to the assumptions of SKAT. The results of the linear model are shown for diverse settings, including varying effect sizes ($\beta$), number of causal variants (CV), whether the effect sizes depend on allele frequency (AF dependent), and whether the allele frequency of causal variants is limited to 5% (max 5%). (**c**) A linear model as in (b), with binary rather than continuous phenotypes, simulated using a step function as in (a). See Supplementary Methods.

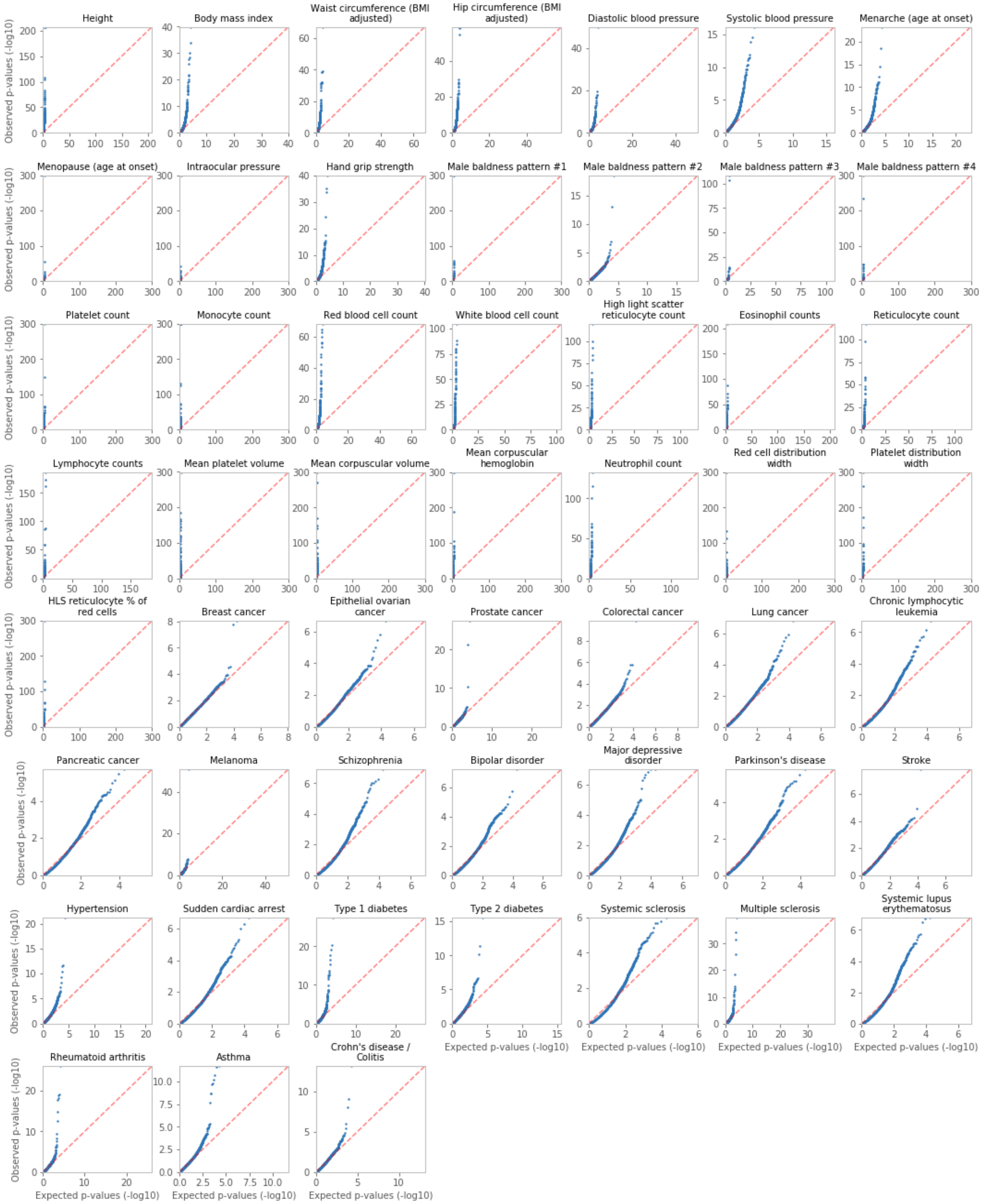


**Supplementary Fig. S2: PWAS QQ plots of all phenotypes**

QQ plots for the obtained PWAS p-values of all 18,053 tested genes, across all 49 tested phenotypes.

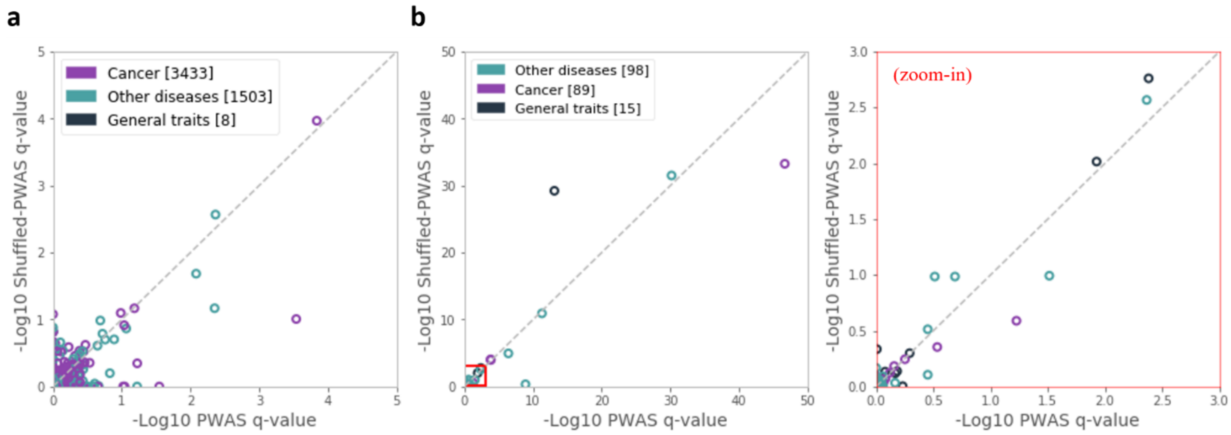


**Supplementary Fig. S3: PWAS with shuffled effect scores**

Comparison between PWAS with the original variant effect scores to an alternative version (named Shuffled-PWAS) where the variant effect scores were randomly shuffled within genes (see Supplementary Methods). (**a**) Comparison between PWAS to Shuffled-PWAS over the 4,944 gene-phenotype associations with strong support by Open Target Platform (as in Fig. 6c). Shuffled-PWAS recovers only 3 of the 6 associations recovered by PWAS ($q < 0.05$), and doesn't recover any additional association. (**b**) Comparison between PWAS to Shuffled-PWAS over the 202 associations reported by OMIM to have a known molecular basis (as in Fig. 6d). Shuffled-PWAS recovers only 10 of the 12 associations recovered by PWAS, and doesn't recover any additional association.

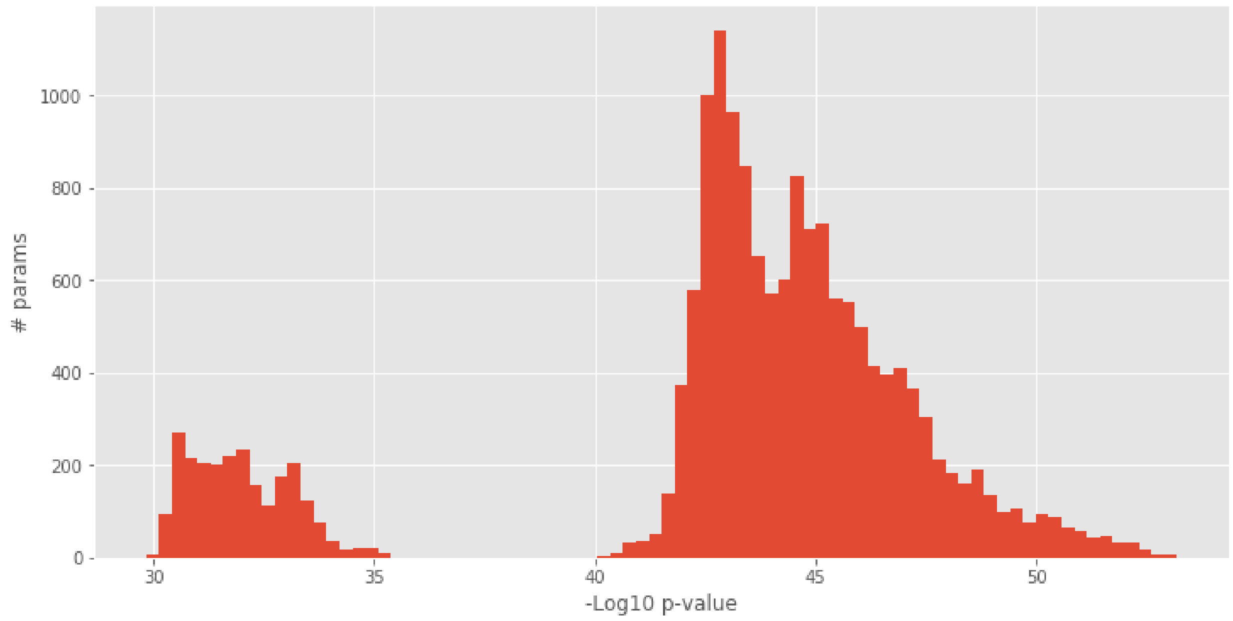


**Supplementary Fig. S4: Distribution of combined p-values over the explored parameters of the effect score aggregation models**

Distribution of the combined p-values (obtained by Fisher's method over the 110 gene-disease OMIM associations used to fit the parameters of the effect score aggregation models) over the 16,807 explored parameter combinations (see Supplementary Methods).

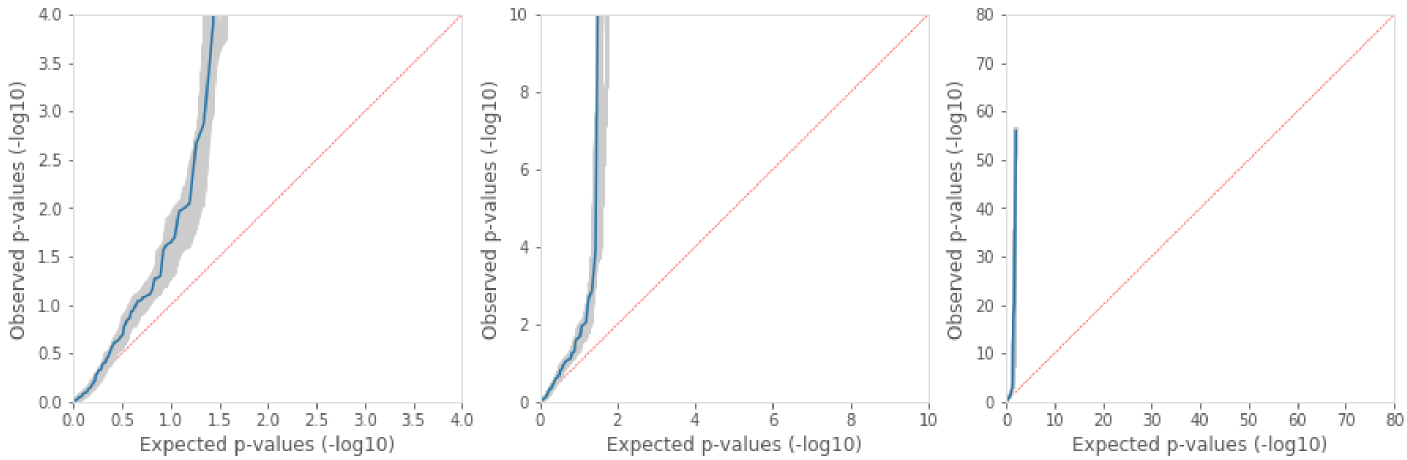


**Supplementary Fig. S5: QQ plot of the explored parameters of the effect score aggregation models**

QQ plots of the obtained p-values for the 110 gene-disease OMIM associations over the explored parameter combinations (see Supplementary Methods), each shown in gray. The p-values of the chosen parameter combination ($\mu_D = 1$, $p_D = 1.25$, $\mu_R = 0.5$, $p_R = \infty$, $q_R = 3$) is shown in blue. The three panels show the same QQ plot in different resolutions.

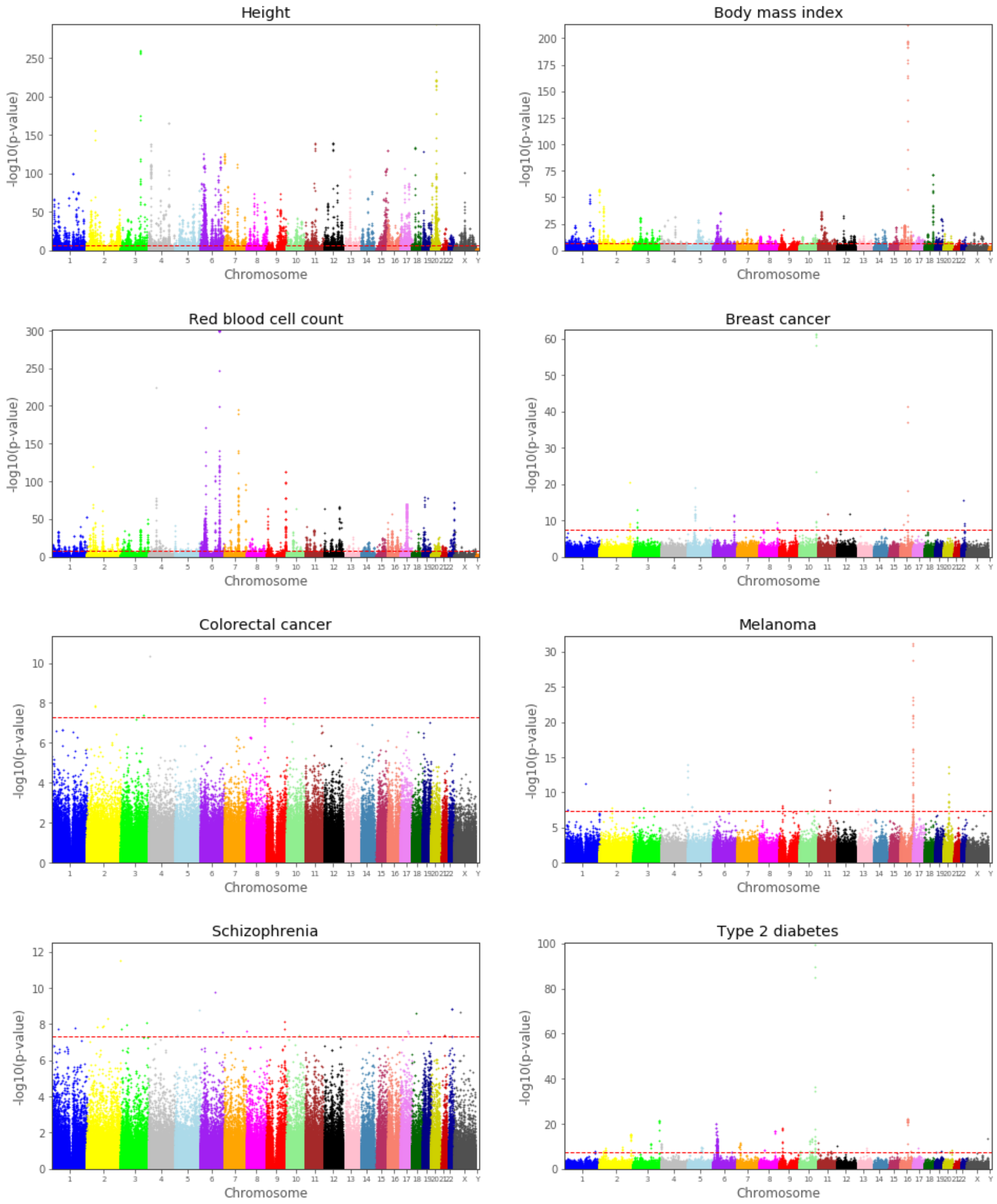


**Supplementary Fig. S6: GWAS Manhattan plots of selected phenotypes**

Manhattan plots, illustrating the GWAS results, for 8 representative phenotypes. High-resolution Manhattan plots for all 49 phenotypes are available online. See Supplementary Methods.

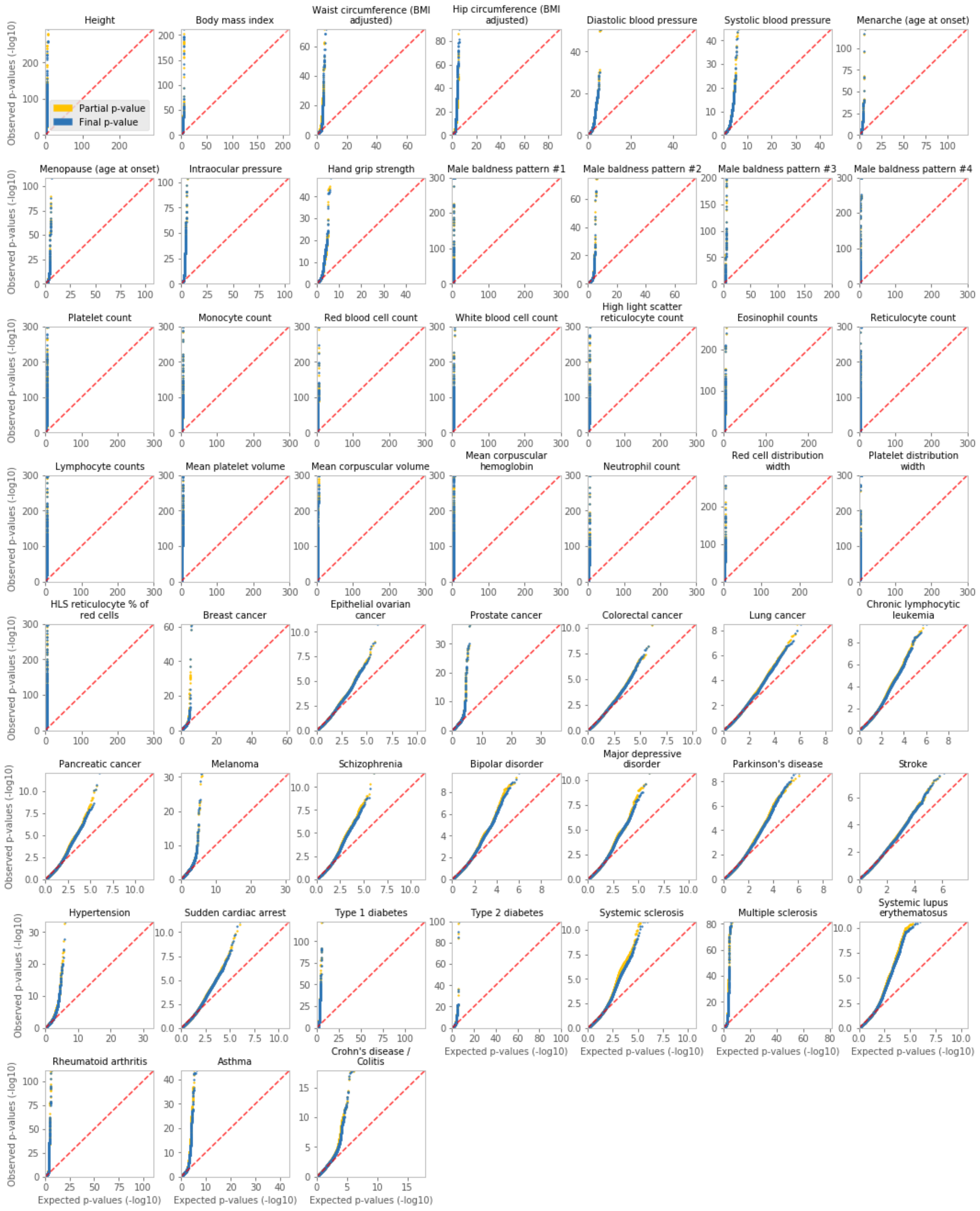


**Supplementary Fig. S7: GWAS QQ plots of all phenotypes**

QQ plots for the GWAS p-values of all tested variants (804,069 raw markers and 639,323 imputed variants), across all phenotypes. Each plot displays the partial p-values (accounting for only a subset of the covariates) and the final p-values (accounting for all covariates, when the partial p-value was below 1E-03). See Supplementary Methods.

### Supplementary methods

#### Parameter fitting of the dominant and recessive effect score aggregation models

The dominant and recessive effect score aggregation models are determined by 5 parameters (2 for the dominant and 3 for the recessive; see Methods). We searched for the combination of parameters that would maximize the discovery power of PWAS. Specifically, we sought to maximize the discovery of accepted and well-supported gene-phenotype associations. To this end, we extracted 110 gene-disease associations from the OMIM database[33] with known molecular basis (mapping key 3). Importantly, the diseases from which we extracted these associations did not overlap with the 49 phenotypes used in the primary analysis reported in this work. The diseases for which all relevant associations were extracted were: psoriasis, gout, agranulocytosis, osteoporosis, deafness, cardiomyopathy (see Supplementary Table S6).

For the $p$ and $q$ parameters, the following 7 values were tried: 1, 1.25, 1.5, 2, 3, 5 and ∞. For the $\mu$ parameter, the following 7 values were tried: 0, 0.1, 0.25, 0.5, 0.75, 0.9, 1. Altogether, 16,807 ($7^5$) combinations were explored. For each combination of parameters $i$ and one of the 110 associations $j$, we applied the generalized PWAS model with that parameter combination and obtained the corresponding p-value $p_{ij}$. For each combination of parameters $i$, we combined the 110 obtained p-values using Fisher's method to obtain a final p-value $p_i$ (Supplementary Fig. S4).We selected the parameter combination that obtained the most significant combined p-value (6.4E-54): $\mu_D = 1$, $p_D = 1.25$, $\mu_R = 0.5$, $p_R = \infty$, $q_R = 3$ (Supplementary Fig. S5).

#### The recessive gene effect score under limit conditions

The recessive effect score of a specific gene in a specific sample is dependent on the values of the $k$ variants affecting the gene: $x_1, \dots, x_k$ and $y_1, \dots, y_k$ (see Methods). Specifically, when $x_i \neq 0$ for all $i \in [k]$, the recessive score is taken to be $R := (1 + \zeta_q) D_p$ where $\zeta_q := \left\| \left( \frac{y_1}{x_1}, \dots, \frac{y_k}{x_k} \right) \right\|_q$ and $D_p$ $:= exp\left( - \left\| \left( \log\frac{1}{x_1}, \dots, \log\frac{1}{x_k} \right) \right\|_p \right)$.

If there exists exactly one $i \in [k]$ such that $x_i = 0$, let us assume without loss of generality that $x_1 = 0$ and $x_2, \dots, x_k > 0$. For conciseness, let us denote $x_1$ and $y_1$ by $x$ and $y$, respectively, and let us also denote $A := \left(\log\frac{1}{x_2}\right)^p + \dots + \left(\log\frac{1}{x_k}\right)^p$ and $B := \left(\frac{y_2}{x_2}\right)^q + \dots + \left(\frac{y_k}{x_k}\right)^q$ (assuming that $p < \infty$ and $q < \infty$, respectively). When $p < \infty$, we can then write $\log D_p(x) = -\left(\left(\log\frac{1}{x}\right)^p + A\right)^{\frac{1}{p}}$.

We are interested in the asymptotic behavior of an expression of the form $(z^p + C)^{\frac{1}{p}}$ when $z \to \infty$ (for some $C \geq 0$ and $p \geq 1$). Let us define $f(t) := (z^p + t)^{\frac{1}{p}}$. Then $f'(t) = \frac{1}{p}(z^p + t)^{\frac{1}{p}-1}$. By the mean-value theorem, there exists $t_0 \in [0, C]$ such that $f(C) - f(0) = (C - 0) \cdot f'(t_0)$, meaning $(z^p + t)^{\frac{1}{p}} - z =$

$C \cdot \frac{1}{p}(z^p + t_0)^{\frac{1}{p}-1}$. Since $\frac{1}{p} - 1 \leq 0$, the function $(z^p + t)^{\frac{1}{p}-1}$ is monotonically decreasing, following that $(z^p + t)^{\frac{1}{p}} - z \leq C \cdot \frac{1}{p}(z^p + 0)^{\frac{1}{p}-1} = \frac{C}{p} \cdot z^{1-p}$. We get that $z \leq (z^p + t)^{\frac{1}{p}} \leq z + \frac{C}{p} \cdot z^{1-p}$. When $p > 1$, we can then write $(z^p + t)^{\frac{1}{p}} = z + o_z(1)$ (with $o_z(1)$ denoting some function satisfying $\lim_{z\to\infty} o_z(1) = 0$).

When $p > 1$, we get that $\log D_p(x) = \log x + o_x(1)$ (with $o_x(1)$ denoting some function satisfying $\lim_{x\to 0} o_x(1) = 0$), resulting $D_p(x) = x \cdot L_1(x)$ with $L_1(x)$ denoting some function satisfying $\lim_{x\to 0} L_1(x) = 1$ (in particular this holds for $p = \infty$ for which $D_p(x) = \min\{x, x_2, \ldots, x_k\} = x$ when $x \to 0$). When $p = 1$, we have $D_p(x) = x \cdot x_2 \cdots x_k$.

Similarly, $\zeta_q(x) = \left(\left(\frac{y}{x}\right)^q + B\right)^{\frac{1}{q}} = \frac{y}{x} + O(1)$ (for all values of $q \geq 1$; in particular for $q = \infty$ for which $\zeta_q(x) = \max\left\{\frac{y_1}{x_1}, \ldots, \frac{y_k}{x_k}\right\} = \frac{y}{x}$ when $x \to 0$).

Overall, we get that $\lim_{x\to 0} R_{p,q}(x) = \begin{cases} y & p > 1 \\ y \cdot x_2 \cdots x_k & p = 1 \end{cases}$. Therefore, we define the recessive model as:

$$R := \begin{cases} 0 & \exists i \neq j, x_i = x_j = 0 \\ y_i & \exists!\, i, x_i = 0, p > 1 \\ y_i \prod_{j\neq i} x_j & \exists!\, i, x_i = 0, p = 1 \\ (1 + \zeta_q) D_p & \forall i, x_i \neq 0 \end{cases}$$

Defining $R$ to be 0 in the case of two distinct variants $i \neq j$ satisfying $x_i = x_j = 0$ makes intuitive sense as $x_i = 0$ represents the event of certain loss of function by variant $i$. Therefore, two certain hits to the gene should indicate a certain recessive effect ($R = 0$).

**Variant effect scores of in-frame indels**

In-frame indels are very rare compared to missense variants[51]. We therefore decided to take a somewhat crude approach for assigning variant effect scores to these variants. Whereas for missense variants we appealed to a rich machine-learning model[22] (which had been trained on the ClinVar dataset[53]), for in-frame indels we ignored the proteomic context of the variants and calculated the effect scores solely based on the numbers of substituted, inserted and deleted amino-acids, according to the formula $s = (1 - P_{sub})^{n_{sub}} \cdot (1 - P_{ins})^{n_{ins}} \cdot (1 - P_{del})^{n_{del}}$, where $n_{sub}$, $n_{ins}$ and $n_{del}$ are the number of amino-acids substituted, inserted and deleted by the complex variant, respectively, and $P_{sub}$, $P_{ins}$ and $P_{del}$ are the empirical frequencies of pathogenic variants among the complex variants (non single-nucleotide) that have resulted in the substitution, insertion or deletion of a single amino-acid, respectively, according to the ClinVar dataset. The obtained values used in this work are: $P_{sub} = 0.71$, $P_{ins} = 0.44$ and $P_{del} = 0.85$.

**Tested protein-coding genes**

In this work, each of the 49 tested phenotypes was tested against 18,053 protein-coding genes for which FIRM provides variant effect scores (in version hg19/GRCh37 of the human reference genome, which is compatible with the genetic data provided by the UKBB). To determine the functional effects of genetic variants on protein-coding genes, FIRM requires multiple layers of genetic and proteomic annotations from a variety of databases (e.g. RefSeq[54], UniProt[49], Pfam[50]). To obtain these annotations, FIRM uses the geneffect Python library (https://github.com/nadavbra/geneffect), constructing 18,053 unified gene objects[22].

**Simulation analysis**

In the simulation analysis (Fig. 3, Supplementary Fig. S1), phenotypes were simulated by the formula $y = \beta x + \sigma$ or $y = \theta(\beta x + \sigma)$ (for continuous or binary phenotypes, respectively), where $x$ is the normalized effect of the gene on the phenotype, $\beta \in \{0.01, 0.05\}$ is the gene's effect size, $\sigma \sim N(0,1)$ is a random Gaussian noise, and $\theta$ is a step function that binarizes the 50% upper values in the cohort into 1 and the 50% lower values into 0. The normalized gene effects $x$ were obtained from the raw gene effects $\tilde{x}$ after subtracting the mean and dividing by the standard deviation of the cohort's samples.

In the protein-centric scheme (Fig. 3, Supplementary Fig. S1a), the gene effects $\tilde{x}$ were simulated according to the framework of PWAS. Specifically, an autosomal protein-coding gene was randomly selected, and the effect scores of all protein-affecting variants were estimated by FIRM. To account for imperfections in FIRM's estimations, a random noise distributed by $N(0, \epsilon)$ was added to the estimated effect scores (for an explored noise parameter $\epsilon \in \{0, 0.25\}$). The resulted variant effect scores were truncated to remain in the $[0,1]$ range. The variant effect scores were then combined with the real UKBB imputed genotypes of the cohort to produce the dominant ($D$) and recessive ($R$) gene effect scores of each sample in the cohort (see Methods). These values were taken to be the gene effect $\tilde{x}$ in the dominant and recessive cases (first and second rows in Fig. 3 and Supplementary Fig. S1a, respectively). In the additive case (third row in those figures), the gene effect was taken to be $\tilde{x} = \frac{1}{2}(D + R)$.

In the linear scheme (Supplementary Fig. S1b-c), the gene effects were simulated as $\tilde{x} = \sum_{i=1}^{m} \beta_i g_i$, where $g_1, \dots, g_m$ are the additive genotypes of $m$ selected causal variants and $\beta_1, \dots, \beta_m$ are random variant effects. The additive genotype of variant $i$ (in the context of a specific sample) is defined as the expected number of alternative alleles, namely $g_i = p_1^{(i)} + 2 \cdot p_2^{(i)}$, where $p_1^{(i)}$ and $p_2^{(i)}$ are the probabilities of 1 or 2 alternative alleles, respectively (these probabilities are the input of imputed genotypes provided by the UKBB). The variant effects $\beta_i$ were selected to be either dependent or independent on allele frequency (denoted "AF dependent" and "AF independent", respectively). In the AF dependent case, the variant effects were chosen by the formula $\beta_i = -\log_{10}(f_i)$ where $f_i$ is the allele frequency of the variant $i$. In the AF independent case, each variant effect was randomly and independently chosen according to a uniform distribution $\beta_i \sim U(0,1)$. When exploring the option of a maximal allele frequency of 5% for causal variants (denoted "max 5%"), the variant effect sizes $\beta_i$ of all variants with an allele frequency $f_i > 0.05$ were set to 0. The number of causal variants was selected to be either $m = 1$ ("one CV"), or all $m = k$ relevant variants ("100% CV"; where $k$ is the number of protein-affecting variants in the selected gene), or 5% of the variants ("5% CV"), i.e. each of the $k$ variants was chosen to be causal or not independently with probability of 0.05.

For each of the three tested methods (GWAS, SKAT and PWAS), we show the statistical power of the method over simulated phenotypes as a function of cohort size. Cohort sizes were taken to be 1,000, 10,000, 50,000, 100,000 or all 332,709 filtered UKBB samples (i.e. the same cohort used in the primary analysis presented in this work; see Methods). Subsets of the full cohort were sampled without replacement. The statistical power of a method (for a specific cohort size and a specific combination of simulation parameters) was estimated as the fraction of simulated phenotypes that obtained a significant p-value with that method. We also present the 95% confidence intervals for these estimations, obtained by approximating the binomial distribution of success or failure as a normal distribution. For each combination of parameters and cohort size, we ran 500 simulations (i.e. 500 independent phenotypes were simulated).

Both PWAS and SKAT were invoked over the same input data (that includes the genotyping of all protein-affecting variants in the selected gene; see the "Running SKAT" section below). For these methods, a simulation result was considered significant if the obtained p-value was below 2.5E-06 (which is the Bonferroni correction of the standard 0.05 threshold, given that 20,000 protein-coding genes are considered). GWAS was considered significant if at least one of the gene's variants obtained a p-value below the accepted exome-wide significance threshold 5E-07. To facilitate feasible computation of p-values by GWAS, we tested only up to 5 variants per simulation, specifically the causal variants with the highest empirical allele frequency in the simulation cohort. In all three methods, all 173 covariates (of the real UKBB data) were included in the calculation of p-values (but they did not play a role in determining the phenotype values). Note that the statistical power of PWAS and SKAT may have been underestimated in the simulation analysis, as FDR correction can be used when they are applied over real data, instead of the more conservative Bonferroni correction used in the simulations.

**Comparable GWAS analysis**

To perform an unbiased comparison between PWAS and GWAS, we applied the two methods on the same dataset derived from the UKBB, and used the same statistical methods (linear and logistic regression, implemented in Python[55]). Across all tested methods we used the same list of 173 covariates (see Methods).

For each phenotype, we tested all 804,069 genetic markers genotyped by the UK Biobank Axiom Array, and 639,323 of the 97,013,422 imputations variants provided by the UKBB[9]. Specifically, we focused only on the 639,323 imputed variants that appeared to affect the coding region of a gene (i.e. the coding region of any of the 18,053 considered protein-coding genes). These are the same 639,323 variants considered by PWAS, meaning that PWAS did not have access to any input data that GWAS didn't. When testing raw genetic markers with GWAS, we used the $g_i \in \{0,1,2\}$ variant encoding (i.e. number of alternative alleles), in order to test for an additive effect (as is the common practice in GWAS). In the case of imputed variants, UKBB provides genotype probabilities rather than definitive called variants, so we took the expected number of alternative alleles, namely $g_i = p_1^{(i)} + 2 \cdot p_2^{(i)}$, where $p_1^{(i)}$ and $p_2^{(i)}$ are the probabilities for 1 or 2 alternative alleles in variant $i$, respectively.

Since running a regression model over ~330K samples with 173 covariates is a computationally-intensive task, we split the analysis of each variant into two stages. First, we ran the regression model only with the sex and year of birth covariates (under the assumption that these are the two most important confounders that need to be accounted for). Only if the p-value obtained in the first stage was lower than 1E-03 we continued to the second stage where we ran the full regression with all 173 covariates. A variant's final p-value was taken to be the one derived from the full regression analysis, if it was run, or from the partial analysis, if it wasn't. A variant was considered significant if its final p-value exceeded the exome-wide threshold (5E-07). In the simulation analysis (see previous section) we did not split GWAS into two stages (i.e. we always performed the regression analysis with all 173 covariates).

The complete GWAS results (of all tested variants against all 49 tested phenotypes) are publicly available at ftp://ftp.cs.huji.ac.il/users/michall/ukbb_gwas_results/. Manhattan plots of the results (merging together the raw markers and tested imputed variants) are shown in Supplementary Fig. S6 for eight selected representative phenotypes. High-resolution Manhattan plots of all 49 phenotypes are publicly available at the same FTP site.

We performed several layers of validation to the GWAS results. To verify that the statistical results behave normally, and that the two-stage approach does not appear to substantially distort the obtained p-values, we examined the QQ plots of both the final and partial p-values (Supplementary Fig. S7). The two-stage approach indeed appears to have only a mild effect on the distribution of p-values. To further validate that the obtained associations are in agreement with established results reported in literature, we performed a comprehensive comparison of our results against the GWAS catalog. Specifically, we compared the enrichment between the significant genomic loci of the two lists (ours vs. the GWAS catalog's results for each phenotype; the mappings of our phenotypes to the GWAS catalog are provided in Supplementary Table S5). For this comparison, we considered different locus radiuses (100, 1K, 10K, 100K and 500K bp). When a certain locus radius was considered, we defined the entire region on the genome whose distance from a significant variant is less than that radius to be significant. If several regions overlapped, we merged them. Following this logic, we obtained a disjoint set of significant genomic regions associated with each phenotype, in each of the two compared resources. We then measured the total length of the genomic regions covered by each of the resources, and the total length of the overlapping genomic segments shared by the two resources. By dividing those values by the total genome length (~3 billion bp) we calculated the fraction of the genome considered to be significant by each of the two methods, and the fraction of the genome considered to be significant by both. Assuming that the two resources are independent, we then calculated the ratio between the observed to the expected overlap fractions, defined as the enrichment factor. In addition to calculating the enrichment factor of all significant associations, we also considered different numbers of maximum loci (100, 1,000 and 10,000). When limiting the number of loci, we took only the top most significant genomic regions (after merging overlapping genomic regions, and considering the p-value of a region to be that of the most significant variant in that region). We then repeated the same process of calculating the enrichment factor, with only the most significant regions considered by each of the two resources. The enrichment factors between our GWAS results to the GWAS catalog (for each of the 49 tested phenotypes, and each of the 20 combinations of parameters) are provided in Supplementary Table S7. Although the approach taken for this comparison is somewhat crude, the enrichments are extremely high. We therefore feel confident that the obtained GWAS results indeed replicate much of the reported associations in the GWAS catalog.

**Running SKAT**

Like with GWAS, we sought to perform an unbiased comparison between PWAS and SKAT[18], by applying the methods on the same UKBB-derived dataset (in particular using the same 173 covariates). We used version 1.3.2.1 of the R package of SKAT (https://cran.r-project.org/web/packages/SKAT/index.html), with 'C' and 'D' output types for continuous and binary/dichotomous phenotypes, respectively.

To allow the two methods access to exactly the same input, we provided SKAT with the genotypes of all protein-effecting imputed variants of each tested gene. Like with GWAS (see previous section), we applied SKAT on additive genotypes (i.e. the expected number of alternative alleles).

**Shuffled PWAS**

To scrutinize the role of the predicted functional effect scores assigned to variants, we compared the performance of PWAS to an alternative version of the method that relies on shuffled effect scores (Supplementary Fig. S3). In the shuffled version of PWAS, within each tested protein-coding gene, the effect scores of all non-synonymous variants affecting the protein sequence (see Methods) were randomly shuffled. We then proceeded with aggregation of the variant effect scores to gene effect scores according to the regular PWAS pipeline (with the same parameters), and significance was calculated by the same statistical analysis (see Methods).

**Summary statistics**

For both PWAS (Supplementary Tables S2-S4) and GWAS (Supplementary Table S1, ftp://ftp.cs.huji.ac.il/users/michall/ukbb_gwas_results/variant_results/), we include, in addition to p-values (and FDR q-values in the case of PWAS), summary statistics that indicate the strength of each association. For continuous phenotypes we include Pearson's correlation coefficient (r), and for binary phenotypes we include Cohen's d (also known as standardized mean difference). Both metrics measure the strength of the correlation between the phenotype values to the genetic values (i.e. the expectation of the number of alternative alleles in the case of GWAS, or the dominant/recessive gene effect scores in the case of PWAS). Note that these metrics do not account for covariates.

It should also be noted that the effect-size metric results end up having opposite interpretations in PWAS and GWAS. In GWAS, a positive value indicates a positive correlation between the phenotype value to the number of alternative alleles, thereby indicating a risk variant. In PWAS, on the other hand, positive values indicate a positive correlation with the gene effect scores, whose higher values mean less functional damage, thereby indicating protective genes. Negative values, on the other hand, have inversed interpretations: protective variants in GWAS vs. risk genes in PWAS.

**Per-phenotype data filtrations**

We constructed the relevant cohort of each phenotype by filtering the 332,709 samples in the main cohort, keeping only the samples with available phenotype values (see Methods).

In binary phenotypes, we sometimes filtered the data to avoid the "dummy variable trap" (i.e. full linear dependence of the variables), as well as complete or quasi-complete separation of the phenotype labels in the logistic regression. Quasi-complete separation occurs when the regression variable values are dependent, conditional on the labels. To avoid such separation (which leads to numeric instability during the training of the logistic regression), we iteratively removed binary covariates that are dependent on the other variables given the phenotype labels, in an order inversed to their frequencies. When removing a binary covariate (e.g. a genetic batch), we also removed all the samples associated with it (e.g. all the samples genotyped in the removed batch), in order to avoid the situation of covariates not accounted for during the statistical analysis.

**Comparison against external resources**

We assessed the quality of discoveries made by PWAS and SKAT by comparing them against two prominent external resources, Open Targets Platform (OTP)[32] and OMIM[33] (Fig. 6).

In Fig. 6b, we calculated the average evidence score of different sets of gene associations (derived by either SKAT, PWAS, or the consensus of the two). To derive the evidence score of a gene (in the context of a given phenotype), we extracted all non-genetic association scores from OTP, i.e. the association scores based on i) affected pathways, ii) animal models, iii) known drugs, iv) literature, v) RNA expression and vi) somatic mutations. We excluded OTP's genetic association score in order to avoid bias by past GWAS discoveries. A gene's evidence score was taken to be the average of the 6 included association scores. To have uniform scale across phenotypes, we normalized the evidence scores into z-values (by taking the mean and standard-deviation of all the ~18K tested protein-coding genes in the context of the same phenotype). The mapping of phenotypes to OTP was made by Experimental Factor Ontology (EFO) accession IDs (Supplementary Table S5). If a given gene symbol had multiple OTP entries, we took the one with the maximum mean non-genetic association score. For each evaluated set of genes, we calculated their average evidence z-value to represent the group (Fig. 6b). In Fig. 6c, we considered all 4,944 gene-disease associations with mean non-genetic association score that was at least 0.1 (in terms of the raw values extracted from OTP, prior to z-value normalization).

In Fig. 6d, we considered the 202 OMIM gene associations implicated with one of the 49 studied phenotypes that have a known molecular basis (mapping key 3). The OMIM entries associated with each phenotype are listed in Supplementary Table S5.

**Criteria of PWAS-exclusive associations**

PWAS-exclusive associations (Table 2, Fig. 7) were defined by the following three criteria: i) very significant (PWAS q-value < 1E-02), ii) without any single dominating variant (the p-value of the most significant nearby variant, up to 500,000 bp to each side of the gene, is not lower than 1E-04), iii) without any other significant gene in the region (up to 500,000 bp to each side of the gene) according to PWAS.

### References


53. Landrum, M. J. *et al.* ClinVar: public archive of interpretations of clinically relevant variants. *Nucleic Acids Res.* **44,** D862--D868 (2015).

54. O'Leary, N. A. *et al.* Reference sequence (RefSeq) database at NCBI: current status, taxonomic expansion, and functional annotation. *Nucleic Acids Res.* **44,** D733--D745 (2015).

55. Seabold, S. & Perktold, J. Statsmodels: Econometric and statistical modeling with python. in *Proceedings of the 9th Python in Science Conference* **57,** 61 (2010).
